# Single cell sequencing as a general variant interpretation assay

**DOI:** 10.1101/2023.12.12.571130

**Authors:** Hongxia Xu, Ling Chen, Mohan Sun, Ken Jean-Baptiste, Dan Cao, Xingyan Zhou, Sandy Wong, Carmen Xiao, Tong Liu, Victor Quijano, Nadav Brandes, Joe Germino, Vasilis Ntranos, Chun Jimmie Ye, François Aguet, Kyle Kai-How Farh

**Author notes:** These authors contributed equally (Hongxia Xu, Ling Chen, Mohan Sun, Ken Jean-Baptiste). These authors contributed equally (Dan Cao, Xingyan Zhou, Sandy Wong). These authors contributed equally (François Aguet, Kyle Kai-How Farh).

## Abstract

The human genome contains ∼70 million possible protein-altering variants, the vast majority of which are of uncertain clinical significance. Closing this gap is essential for accurate diagnosis of disease-causing variants and understanding their mechanisms of action. Towards this goal, we developed a pooled perturbation approach combining saturation mutagenesis with single cell RNA sequencing to map the effects of every single nucleotide variant in a gene. We sequenced ∼440,000 cells expressing variants in *CDKN2A* (p16^INK4a^), *TP53*, and *SOD1*, observing almost all possible protein-coding variants, with a mean of 61 cells per variant. Using single cell gene expression signatures, we show that each gene may contain multiple types of pathogenic variants that affect distinct downstream pathways. We demonstrate that single cell expression signatures outperform existing bulk experimental assays and computational models for predicting pathogenicity, and summarize both the utility and potential limitations of single cell sequencing as a general variant interpretation assay.

## Introduction

Understanding the functional consequences of genetic variation is key to identifying pathogenic variants and their role in disease. Only a small fraction of all possible ∼70 million protein-altering variants in the human genome have been categorized and annotated in clinical variant databases such as ClinVar (Landrum et al., 2016), with the vast majority remaining variants of uncertain significance (VUS). Significant progress in variant characterization has been made using high-throughput saturation mutagenesis experiments also known as deep mutational scanning (Fowler and Fields, 2014), which typically rely on cellular survival or proliferation as a readout of pathogenicity (Findlay et al., 2018; Giacomelli et al., 2018; Matreyek et al., 2018). However, such one-dimensional readouts preclude identifying the functional mechanisms associated with individual variants or distinguishing between different modes of pathogenicity, or the contexts wherein they may arise. Moreover, while similar progress to predict variant pathogenicity has been achieved with deep learning models (Brandes et al., 2023; Cheng et al., 2023; Frazer et al., 2021; Gao et al., 2023), these models also lack a functional readout.

In combination with saturation mutagenesis assays, where each cell receives a different perturbing variant, single cell sequencing has the potential to provide a rich expression signature for each individual perturbation (Ursu et al., 2022). Although RNA-seq from individual single cells is sparse, averaging expression across cells that received the same variant can provide robust gene expression signatures of the perturbed downstream pathways, while also minimizing batch effects between perturbations (Replogle et al., 2022). Recent Perturb-seq experiments with CRISPRi and CRISPRa on the scale of thousands of genes have also shown that single cell expression signatures can serve as a general phenotypic readout (Replogle et al., 2022; Schmidt et al., 2022), reducing the necessity of developing bespoke assays for individual genes, which may only measure a subset of their functions. To date, single cell expression readouts have only been performed for a limited number of variants per gene (Ursu et al., 2022), nevertheless showing that the effects of benign and pathogenic variants can be distinguished by their transcriptomic signatures. This existing method relied on fixed molecular barcodes to encode the identity of 100 variants, represented only a small fraction (∼3%) of the total number of possible exonic single nucleotide variants (SNVs). To construct saturation mutagenesis libraries containing the thousands of possible variants in most genes, we instead use random barcodes to encode each variant and read out the variant corresponding to each barcode in a separate sequencing step.

We apply this approach to characterize almost all amino acid variants encoded by SNVs in three genes: *CDKN2A* (cyclin-dependent kinase inhibitor 2A), specifically the p16^INK4a^ isoform, a tumor suppressor and cell cycle regulator that is inactivated by somatic mutation in many cancers and for which germline mutations underlie familial melanoma (Hussussian et al., 1994) and pancreatic cancer (Goldstein et al., 1995); *TP53* (the tumor suppressor p53), the most frequently mutated gene across human cancers wherein germline mutations result in the cancer-predisposing Li-Fraumeni syndrome (McBride et al., 2014); and *SOD1* (superoxide dismutase 1), the first gene causally linked to amyotrophic lateral sclerosis (ALS) (Rosen et al., 1993), with SOD1 mutations accounting for ∼12% of familial and ∼1-2% of sporadic ALS cases (Goutman et al., 2022). Using gene expression profiles from hundreds of thousands of cells expressing mutant alleles of these genes, we apply machine learning approaches to identify distinct modes of pathogenicity linked to specific structure-function relationships. We benchmark our pathogenicity scores against computational predictions and prior deep mutational scanning experiments to demonstrate that our single cell assay enables state-of-the-art pathogenicity prediction coupled to a functional interpretation of variant effects.

## Results

### Saturation mutagenesis with single cell RNA sequencing readout

We generated saturation mutagenesis libraries for the three genes (Methods), obtaining close to uniform coverage for all possible SNVs in *TP53* and *SOD1*, but with some regions of low or missing coverage in *CDKN2A* due to synthesis limitations related to high GC content (Figure S1A). The libraries were cloned into lentiviral vectors containing in the 3’ UTR a 19 bp semi-random variant barcode designed to minimize homopolymer and polyadenylation motif formation (Methods), resulting in over ∼750,000 different variant barcode–variant combinations per gene (Figure S1B). For each gene, these barcoded lentiviral vectors were then transduced into cells at low multiplicity of infection (MOI; Methods and Figure 1) to target a single variant per cell. All experiments were conducted in A549 lung carcinoma epithelial cells, which have a homozygous deletion of the *CDKN2A* locus (Ikediobi et al., 2006). Following lentiviral transduction, we prepared 3’ scRNA-seq libraries using the 10x Chromium protocol, from which the variant barcode(s) in each cell were read out directly (Figure 1 and Methods).

**Figure 1.**
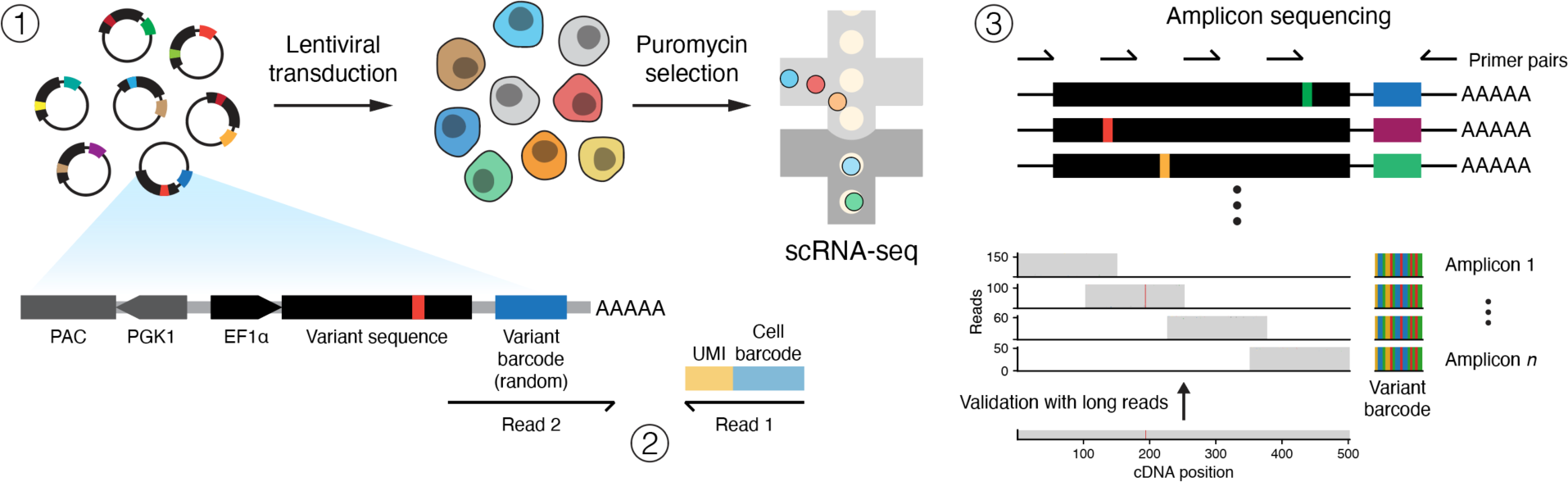
Saturation mutagenesis with single cell RNA-seq readout. Schematic representation of the experimental workflow, consisting of 1) lentiviral transduction of a saturation mutagenesis library with random variant barcodes, followed by single cell RNA sequencing (scRNA-seq); 2) linking of variant barcodes to their corresponding cell barcode directly from the scRNA-seq data; and 3) targeted amplicon sequencing of the single cell cDNA library with a tiling set of primers, linking each random variant barcode to a specific mutation. Bottom panels: read pileups from four amplicons of a variant transcript in the TP53 library, with corresponding long read data for validation.

To recover the variant corresponding to each variant barcode, we performed amplicon sequencing on the cDNA library from each experiment, with primers designed to amplify exogenous transcripts and tile the coding sequence of each gene (Figures 1 and S1C and Methods). Because lentiviral recombination can rearrange variant barcode–variant pairings, we performed amplicon sequencing on each experiment to maximize the accuracy of variant assignments. We developed a custom pipeline to call variants from amplicon sequencing, using stringent quality controls to only retain barcodes that could be confidently assigned to a single variant or wild type (WT) sequence (∼1-22% of each library consisted of WT transcripts; Figure S1B and Methods). To verify the accuracy of our variant calls, we sequenced the TP53 library using long reads, obtaining >99% concordance (Figures 1, S1C and S1D and Methods).

We integrated the variant calls with the scRNA-seq data by matching on variant barcodes, retaining only cells that contained a single variant and following standard quality control procedures (Luecken and Theis, 2019) to select cells for analysis, from which we generated the covariate-corrected and normalized gene expression matrices (as z-scores; Methods) that we used to characterize variant effects on the transcriptome.

### A gradient of pathogenicity of p16 variants linked to disruption of CDK6 binding

After applying quality controls, we obtained 141,973 single cells expressing 1,389 of 1,413 (98.3%) possible CDKN2A/p16^INK4a^ (henceforth p16) variants, with 1,230 SNVs (88.6%) observed in at least 20 cells, corresponding to 964 of 1,069 (90.2%) possible amino acid variants (Figure S2A). We excluded 81 SNVs due to insufficient coverage in the saturation mutagenesis library (Figure S1A and Methods). Assigning each cell to a cell cycle phase based on its expression profile showed that cycling cells were enriched for PrimateAI-3D computationally-predicted pathogenic variants (Gao et al., 2023) (Figures 2A, 2B and S2B and Methods), consistent with the role of p16 in inhibiting cell proliferation by binding to cyclin-dependent kinases 4 and 6 (CDK4/6) (Russo et al., 1998). For simplicity, we primarily reference the PrimateAI-3D algorithm in this work given its previously documented performance on a wide range of benchmarks (Gao et al., 2023) and because it did not use human-annotated clinical variant databases for training, while also providing comprehensive benchmarking for other computational methods (Methods). Averaging the expression profiles across cells for each variant revealed a separation into two unsupervised clusters (Methods) corresponding to likely pathogenic and synonymous-like benign variants, respectively, on the basis of PrimateAI-3D scores (Figure 2C) and ClinVar annotations (Figure 2D).

**Figure 2.**
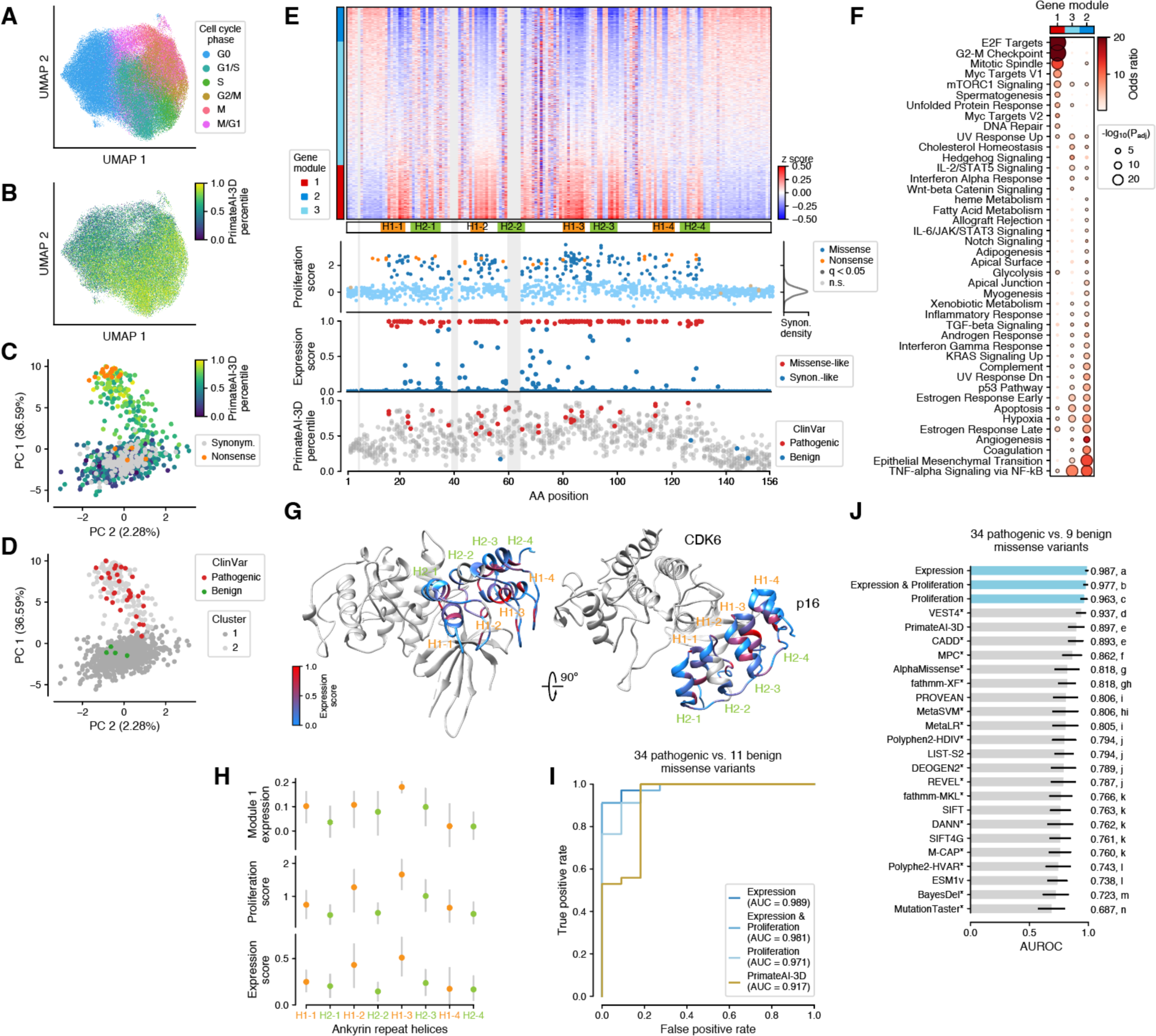
Saturation mutagenesis of p16 variants with single cell RNA-seq readout. (A) UMAP of expression profiles for 141,973 single cells in A549 cell culture, each expressing a p16 variant in a null background, labeled by cell cycle phase (Methods). (B) Same as (A), with missense variants labeled by PrimateAI-3D score, showing a correlation between variant pathogenicity and cycling cells. (C) PCA of expression profiles for amino acid variants detected in ≥30 cells, with missense variants labeled by PrimateAI-3D score. (D) Same as (C), with ClinVar variants labeled (Methods). (E) Top: heatmap showing the average expression profile (z-scores) at each residue position (Methods). Bottom panels: proliferation score for protein-altering variants, with significantly enriched variants labeled (q-value < 0.05; Methods); expression score for missense variants (Methods); PrimateAI-3D scores for all missense variants, with ClinVar variants labeled. Light gray: regions excluded due to low coverage in the saturation mutagenesis library. (F) Biological pathways enriched for gene modules up- or down-regulated upon expression of p16 variants. (G) Structure of p16 bound to CDK6 (PDB: 1BI7), with p16 residues labeled by expression score. (H) Expression levels of module 1 genes, proliferation score, and expression score at residue positions within the helices comprising the ankyrin repeats of p16, showing that variants in helix 1 are on average more pathogenic. Errors bars indicate 95% confidence intervals. (I) ROC curves and area under the curve (AUC) for single cell–derived scores (expression, proliferation, and expression & proliferation ensemble scores), and PrimateAI-3D predictions for distinguishing ClinVar pathogenic from benign variants. (J) Classification results for 22 computational prediction algorithms on the subset of variants from (I) scored by all methods, with methods that were exposed to human clinical variant annotations or allele frequency databases during training labeled with an asterisk to indicate a risk of overfitting. Error bars: standard deviations from bootstrapping.

We trained a classifier on per-variant averaged gene expression profiles to distinguish between synonymous and nonsynonymous variants (Methods), defining pathogenicity as the probability of a variant being nonsynonymous; we refer to this as the ‘expression score’ (Figure 2E). Using this score, we classified 155/898 missense variants and 17/22 nonsense variants as pathogenic (Methods and Table S1). We also assigned a ‘proliferation score’ to each variant, calculated as the number of cells expressing the variant normalized to its abundance in the original library (Methods). The two scores were in close agreement, with 96.4% of missense and all nonsense variants assigned the same label by both (Figures 2E and S2C), and were well-correlated with PrimateAI-3D predictions (Figure S2D; Spearman’s ρ = 0.54-0.58). The proliferation score also replicated findings from a prior study that measured growth rates for a limited number of p16 variants (n=43, ∼4% of all variants) in pancreatic ductal adenocarcinoma cell lines (Kimura et al., 2022) (Spearman’s ρ = 0.74-0.78, Figure S2E).

To identify relationships between variant effects and protein structure, we computed average gene expression profiles at each amino acid position in the protein, weighting the profiles of different missense variants by their expression score (Figure 2E). This showed that pathogenic variants were primarily concentrated in the four ankyrin repeats of p16, each consisting of two alpha helices, H1 and H2 (Figure 2E), as well as in the loops connecting each pair of helices, with variants in the N and C termini appearing to be benign. Pathogenic variants displayed robust downstream transcriptomic signatures, including upregulation of E2F target and G2-M checkpoint genes (gene module 1) and downregulation of genes in pathways including TNFα signaling, epithelial to mesenchymal transition, and apoptosis (Gil and Peters, 2006) (gene modules 2 and 3; Figures 2E and 2F). Most nonsense variants were also pathogenic as expected given the *CDKN2A* null background, except for variants near the end of the protein, which had synonymous-like effects in our assay. The p.Glu2Ter nonsense variant also displayed a synonymous-like effect, consistent with stop codon read-through or translational re-initiation (Dabrowski et al., 2015).

Mapping the expression score onto the structure of p16 bound to CDK6 (Russo et al., 1998) further showed that the residue positions with the highest average pathogenicity were those closest to the binding interface between the two proteins, as well as internal residues (Figure 2G). Variants in the H1-2 and H1-3 helices produced the largest effects (Figure 2H), consistent with their role in binding CDK6 (Russo et al., 1998). In contrast, in the H2 helices, which do not interface with CDK6, the outward-facing residue positions were more benign than the inward-facing ones. These results were consistent across multiple measures of variant effects, with the activation of gene expression in the module 1 pathway and our expression and proliferation scores all implicating the residues in the H1 helices in direct contact with CDK6 (Figure 2H).

Next, we sought to benchmark our pathogenicity scores on the task of classifying pathogenic from benign variants from the ClinVar clinical variant database. Since only four p16 variants were annotated as benign in ClinVar, we also included 7 variants observed at allele frequencies > 0.00005 across gnomAD, TOPMed, and UK Biobank for benchmarking purposes, reasoning that such variants were likely benign. We chose this threshold to maximize the number of putatively benign variants, while verifying that the findings were robust across a range of allele frequency thresholds (Figure S2F). We benchmarked our scores on a common set of 34 pathogenic and 9 putatively benign variants against predictions made by 22 computational models (Methods), including SIFT (Ng, 2003), PolyPhen-2 (Adzhubei et al., 2013), REVEL (Ioannidis et al., 2016), CADD (Rentzsch et al., 2019), AlphaMissense (Cheng et al., 2023) and PrimateAI-3D; this included methods that were directly trained on human variants, such as clinical variant annotations (e.g., ClinVar, HGMD) and human allele frequency databases, as well as methods trained without such annotations (Methods). Our expression score achieved the best classification performance (ROC AUC = 0.987), followed by the proliferation score (ROC AUC = 0.963, Mann-Whitney P = 3.1e-85), with all 22 computational predictions performing worse, despite some of these methods having trained on annotated variants that overlapped the benchmarking set (Figures 2I, S2G and 2J). When restricting the benchmarking set to variants that were observed in at least 30 cells in our assay, the expression score perfectly classified pathogenic and benign variants (AUC = 1.0; Figure S2H), indicating that the one discordant missense variant was likely misclassified due to insufficient coverage (13 cells compared to an overall average of 85.7 cells per variant). Based on these results, the expression score enabled classification of 47 ClinVar VUS as likely pathogenic and 317 as likely benign (with 4 not scored due to insufficient coverage). The lower performance of computational predictors in this benchmark likely relates to pathogenic p16 variants disrupting specific binding interactions with CDK4/6 and insufficient protein-protein contacts used to train these models. Taken together, our results show that single cell gene expression profiles accurately capture perturbations induced by pathogenic variants and enable state-of-the-art prediction of variant pathogenicity in p16.

### Domain-specific pathogenic effects of TP53 missense variants

We sequenced 206,394 single cells expressing TP53 variants on a A549 wild-type background, observing all 3,546 possible SNVs, with 3,442 (97.1%) in at least 20 cells, corresponding to 2,739 of possible 2,814 (97.3%) amino acid variants (Figure S3A). Similar to p16, unsupervised two-dimensional projection of single cell gene expression profiles showed that cycling cells were enriched for computationally-predicted pathogenic variants (Figures 3A, 3B and S3B), consistent with the role of TP53 in regulating cell cycle progression (Hafner et al., 2019) and dominant negative effects of TP53 missense mutations (Boettcher et al., 2019; Giacomelli et al., 2018). While this association was relatively weak for individual cells, averaging the expression for each variant revealed distinct clusters of variants with synonymous-like and nonsynonymous-like expression profiles (Figures 3C and S3C).

**Figure 3.**
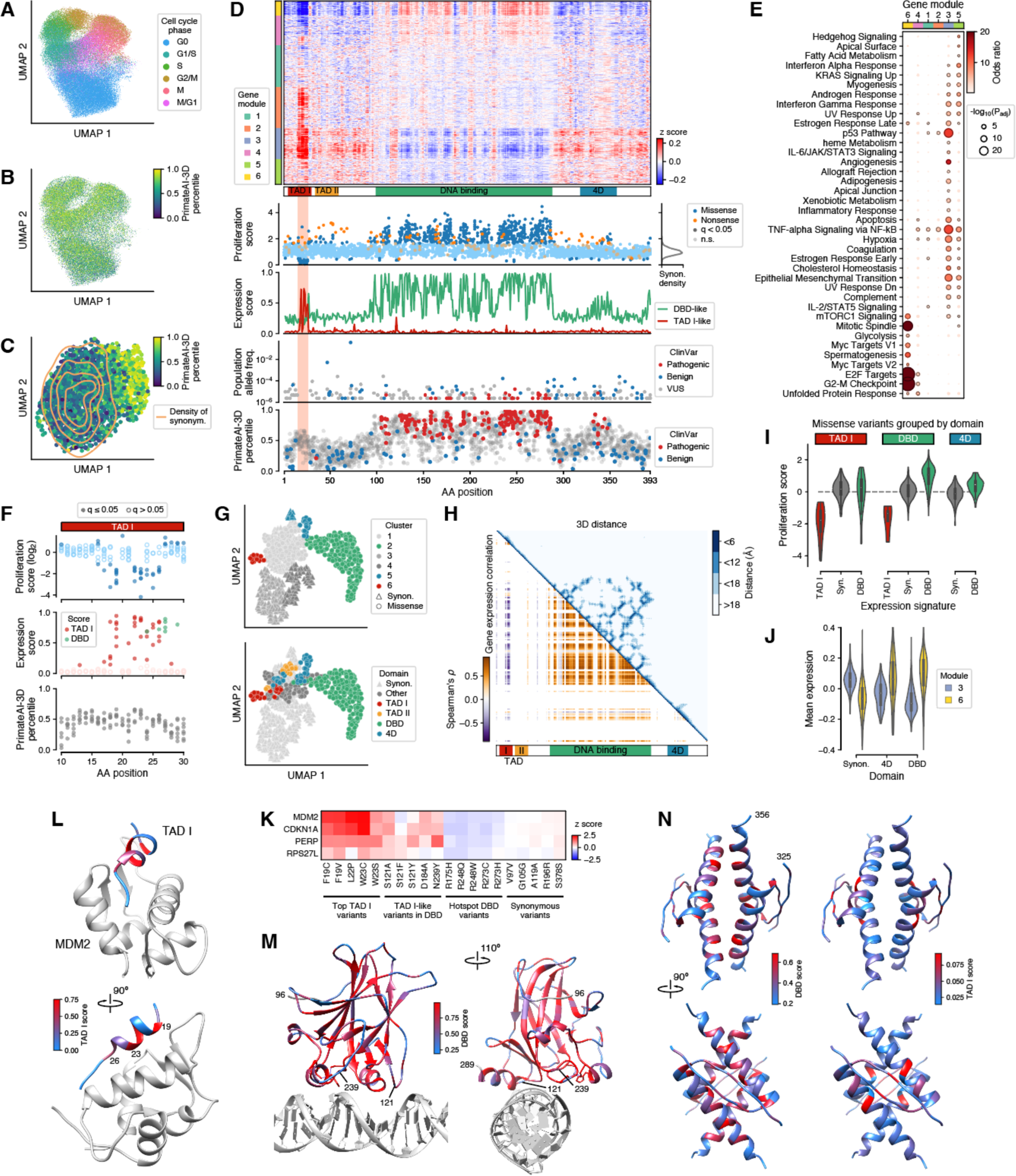
Domain-specific variant effects in TP53 are associated with multiple axes of pathogenicity. (A) UMAP of expression profiles from 206,394 single cells in A549 cell culture, each expressing a TP53 variant on a wild-type background, labeled by cell cycle phase (Methods). (B) Same UMAP as in (A), showing missense variants labeled by PrimateAI-3D. (C) UMAP of expression profiles for missense variants labeled with PrimateAI-3D scores, with the distribution of synonymous variants shown as density contours. (D) Top: heatmap showing the expression profile (z-score) averaged across neighboring residue positions (Methods). Bottom panels: proliferation score for protein-altering variants, with significantly enriched variants labeled (q-value < 0.05; Methods); pathogenicity scores (expression score; Methods) for variants with DBD-like or TAD I–like expression signatures; allele frequency of germline TP53 variants across ∼416,800 individuals, showing an absence of variants in a region of TAD I (highlighted in red across panels); PrimateAI-3D scores for all missense variants, with ClinVar variants labeled. (E) Biological pathways enriched for gene modules up- or down-regulated upon expression of TP53 variants (Methods). (F) Experimental scores and PrimateAI-3D predicted pathogenicity in TAD I. (G) UMAP of expression profiles averaged across neighboring residue positions (Methods), labeled by unsupervised clusters (top) and domain annotations (bottom). (H) Correlation between the average expression profiles for pathogenic variants at each residue position, juxtaposed with the contact map from the AlphaFold prediction. (I) Proliferation scores for variants classified as having TAD I-, DBD-, or synonymous-like expression signatures across domains. (J) Mean expression of genes in modules 3 and 6 for variants classified as having TAD I-, DBD-, or synonymous-like expression signatures. (K) Expression of core p53 pathway genes for variants with TAD I-, DBD- and synonymous-like signatures. (L-N) Structures of MDM2 bound to TAD I (L; PDB: 1YCR), DBD (M; PDB: 1TUP) and 4D tetramer (N; PDB: 1C26), labeled by TAD I- and DBD-like expression scores.

To globally identify the effects of variants on gene expression and their functional mechanisms, we next computed average gene expression profiles at each amino acid position in the protein (Figure 3D). Since the average expression changes were weaker compared with p16 variants due to the expression of TP53 variants on a wild-type background, we additionally averaged neighboring residue positions in the 3D protein structure (Methods). The resulting expression profiles revealed distinct, domain-specific signatures of variants, notably in Transactivation Domain I (TAD I) and the DNA-binding domain (DBD) (Figure 3D). We identified six distinct gene modules by unsupervised clustering of gene expression (Methods), including upregulation of E2F target and G2-M checkpoint genes for variants in the DBD (module 6), and an apoptosis signature (module 4) and a p53/TNFα signaling signature (module 2) for variants in TAD I (Figure 3E). These changes in gene expression were consistent with effects on proliferation, with the highest proliferation scores (computed as for p16; Methods) observed for DBD variants, whereas TAD I variants, which produce an apoptosis signature (module 4), led to a depletion of cells (Figures 3D and 3F). Apoptosis-inducing TAD I variants were not observed in a population of ∼416,800 individuals (aggregated across gnomAD, TOPMed, and UK Biobank; Figure 3D), consistent with strongly apoptotic effects being incompatible with survival.

Two-dimensional projection and unsupervised clustering of gene expression profiles (Methods) grouped residue positions by domain, with effects of variants in TAD I, DBD, and the tetramerization domain (4D) being distinct from any other domain (Figures 3G and S3D). In contrast, variants in Transactivation Domain II (TAD II), as well as variants in unstructured regions, did not produce expression signatures that were distinct from synonymous variants (Figure 3G). Based on these results, we trained a classifier to differentiate between TAD I-, DBD-, 4D- and synonymous-like signatures (Figures 3D and S3E, and Table S2). We found that variants in the 4D domain were classified similarly as DBD variants, but with weaker effects on proliferation (Figure 3I) and gene expression, with lower average down- or up-regulation of module 3 and 6 genes, respectively (Figure 3J). In addition, variants at specific positions within the DBD domain, notably at S121 and N239, displayed different signatures from their neighbors and were classified as TAD I-like variants by their expression profiles, as well as having negative effects on cell proliferation (Figure 3D). When we computed the gene expression correlation matrix for all variants with non-synonymous expression signatures, residue positions with correlated gene expression signatures corresponded to physical contacts in the 3D structure (Figure 3H), reflecting the two major axes of pathogenicity around the DBD and TAD I domains. The classifier included predictions for nonsense variants (Figure S3F), and we found that these generally had DBD-like effects, except when the variant occurred before the TAD II domain or at the C terminus, in which case they had synonymous-like expression signatures. These findings were consistent with previously published results from a bulk saturation mutagenesis assay performed in a TP53 null background (Giacomelli et al., 2018) (Figure S3F), although these experiments should be cautiously interpreted in the context of non-physiological nonsense-mediated decay in exogenous transcripts.

Next, we sought to characterize in detail the structure-function relationships underlying domain-specific pathogenic effects, starting with specific amino acid positions in TAD I. The hydrophobic residues F19, L22, W23, L25 and L26 had high TAD I-like expression scores (Figures 3F) and are deeply conserved and essential for binding to other proteins (Raj and Attardi, 2017), including MDM2, which negatively regulates TP53 by blocking transactivation. Indeed, mapping the average TAD I–like expression score onto the structure of TAD I bound to MDM2 showed that residues at the interface between the two proteins had the largest variant effects (Figure 3L), consistent with disruption of MDM2 binding leading to p53-induced apoptosis (Chène, 2003). In contrast, residue P27, which is also deeply conserved and is essential for binding by the negative regulator adenoviral oncoprotein E1B, had a low average TAD I-like score, a high DBD-like score, and increased proliferation (Figure 3F). The P27Y mutation has previously been shown to disrupt E1B binding while increasing MDM2 binding affinity (Lin et al., 1994), leading to MDM2 sequestration and increased proliferation.

In the DBD domain, a majority of missense variants (681/1117, 61%) had high DBD-like expression scores, with the strongest effects localized predominantly in the core of the DNA binding domain (Figure 3M). The four variants with outlier TAD I–like signatures, S121F, S121A, S121Y and N239Y were at the periphery, towards the DNA binding interface (Figure 3M). The variants at S121 have been previously characterized as ‘super’ mutants leading to increased apoptosis induction (Kakudo et al., 2005; Saller, 1999), with S121F exhibiting differential specificity in activating the MDM2 promoter compared to G-rich promoters (Saller, 1999), whereas N239Y was shown to increase the stability of the DNA-binding domain (Nikolova et al., 1998). Indeed, in our experiments, we observed lower MDM2 expression in cells expressing S121F (Figure 3K), whereas cells expressing N239Y instead showed increased levels of PERP, a downstream target of TP53 that promotes apoptosis (Attardi et al., 2000).

To gain insights into the functional effects of 4D variants, we mapped both the DBD- and TAD I–like expression scores onto the 4D tetramer assembly (Figure 3N). Residues with high average DBD-like expression scores were predominantly located at the interface of the four domains, with few outward-facing mutations having pathogenic effects. Conversely, residues with higher TAD I-like scores were specific to the antiparallel beta sheet and helix-helix interfaces between 4D dimers (Figure 3N). These observations were consistent with prior studies of individual variant effects on p53 oligomerization (Fischer et al., 2016). Overall, variants in the tetramerization domain had weaker effects than variants in the DBD and TAD I domains, although they perturb similar transcriptional programs.

Finally, variants at residues D391, S392, and D393 at the C terminus, which are deeply conserved (Laptenko et al., 2016), produced weakly elevated DBD-like expression and proliferation scores, similar to variants in 4D (Figure 3D). Phosphorylation of S392 has been shown to increase tetramer formation (Sakaguchi et al., 1997), and mutations at this residue would therefore likely mimic 4D mutations, as observed in our experiments.

Taken together, our results reveal two major axes of pathogenicity in TP53, with either TAD I–like apoptotic effects or DBD-like effects on cell proliferation. While these effects apply to most variants in these domains, the scRNA-seq readout enabled the identification of individual variants within each domain that are clearly distinct in the downstream molecular pathways that they activate.

### TP53 variant classification performance

The characterization of distinct modes of pathogenicity in TP53, combined with the relatively large number of TP53 missense variants annotated as either pathogenic or benign in ClinVar (189 and 107, respectively) presented an opportunity to comprehensively benchmark the performance of our single cell assay against prior bulk mutational scanning experiments (Giacomelli et al., 2018) and 27 published computational prediction algorithms (Methods), on DBD- and TAD I–like pathogenicity. On the task of classifying ClinVar pathogenic vs. benign variants, our ensemble classifier based on proliferation and expression (i.e., having either pathogenic DBD- or TAD I–like effects) achieved the best performance (ROC AUC = 0.983), followed by the computational methods, which differed widely in performance (Figures 4A and 4B). We flagged the computational methods that used clinical variant annotations or human allele frequency databases during training and validation to caution that their performance in the benchmark may be inflated due to overfitting. Our single cell–derived scores also achieved superior performance compared with the proliferation-based bulk assay from (Giacomelli et al., 2018), which was conducted under similar experimental conditions (i.e., overexpression of mutant TP53 alleles in A549 cells with a wild-type background).

**Figure 4.**
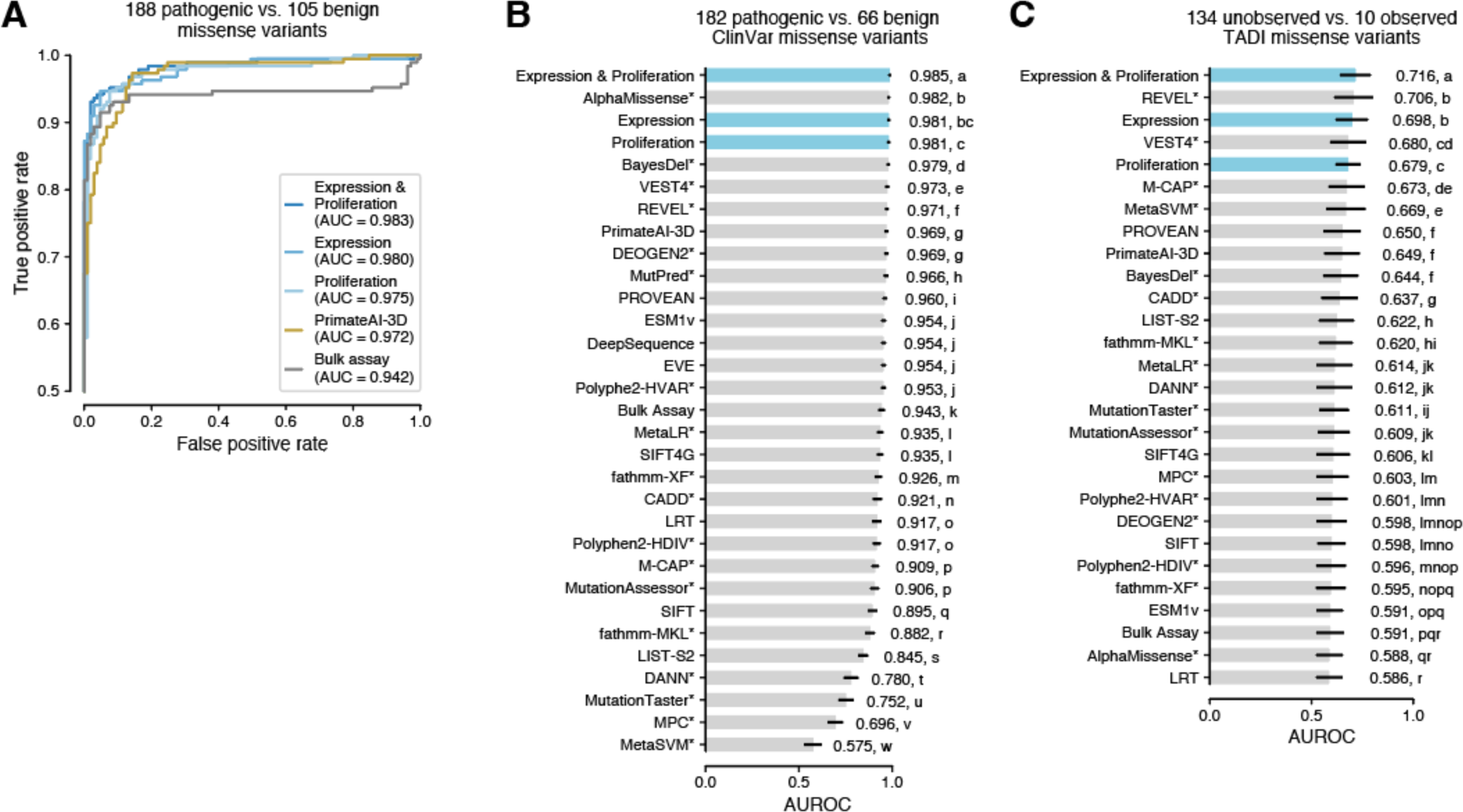
Variant classification performance for TP53. Benchmarking of single cell–derived scores (expression alone (Expression), proliferation alone (Proliferation), and ensemble of the two scores (Expression & Proliferation)) against computational predictors of pathogenicity (Methods) on the task of distinguishing pathogenic from benign variants annotated in ClinVar. (A) ROC curves and area under the curve (AUC) for the expression, proliferation and ensemble scores, the bulk assay dominant-negative score from (Giacomelli et al., 2018), and PrimateAI-3D predictions. (B) Performance of experimentally derived scores and computational models on the task of classifying pathogenic vs. benign ClinVar variants, with models that were trained using ClinVar or other clinical variant annotations or human allele frequency databases indicated by *, and scores from this work labeled in blue. Error bars indicate standard deviations of the scores from bootstrapping (Methods). (C) Comparison of computational methods on the task of distinguishing between benign TAD I variants that were observed in the population vs. variants that have never been observed.

Next, we sought to compare classification performance for variants in TAD I. Earlier, we observed that variants found in human populations are strongly depleted in this domain, consistent with apoptotic effects being incompatible with survival. As neither pathogenic or benign ClinVar variants are annotated in this domain, we benchmarked performance on the task of distinguishing between variants observed in at least one person in the population (10) vs. variants that were never observed in ∼416,800 individuals from the gnomAD, TOPMed, and UK Biobank cohorts (134). Although classification performance was lower compared to the ClinVar benchmark, our ensemble classifier also achieved the highest performance on this task (ROC AUC = 0.716; Figure 4C). Similar to our earlier observations for p16, the computational models generally showed weaker performance when predicting the pathogenicity of variants that affect protein-protein interactions.

While we observed strong agreement between the ensemble score from our assay and the 295 missense variants annotated in ClinVar, there were a few discrepancies (6%), including ten ClinVar pathogenic variants that were scored as benign by our ensemble classifier. Nine of these variants were also scored as benign in the bulk assay from (Giacomelli et al., 2018), raising the possibility that they may not have dominant negative pathogenic effects. Among them, H296R was predicted least likely to have a pathogenic effect (ensemble score = 0.086). This variant was only observed in one individual in COSMIC, further supporting an absence of dominant negative effects, and seven other amino acid variants at the same position were also classified as benign by both our assay and the bulk assay from (Giacomelli et al., 2018), consistent with low overall pathogenicity of variants at this position. The remaining variant, Q165L, was classified as borderline benign by the ensemble score (0.365; FDR = 0.105) and may thus have intermediate effects. Conversely, eight variants annotated as likely benign in ClinVar were scored as pathogenic by our ensemble classifier. These variants had intermediate PrimateAI-3D percentile scores ranging from 0.25-0.76, and five were at amino acid positions with other variants with pathogenic effects in our assay. For R110H, two other variants at the same position (R110L and R110P) were annotated as pathogenic in ClinVar. Together, this suggests likely intermediate effects or pathogenicity of these variants. One variant (S303G) had coverage below 99.55% of all cDNA variants in the library, potentially producing a noisy estimate of its effects. The remaining variant, D393Y, which was confidently classified as a pathogenic variant by our ensemble classifier (0.844, FDR 0.008) was not observed in the germline genomes of ∼416,800 individuals and occurred at the deeply conserved D393 residue, indicating its likely pathogenicity (Laptenko et al., 2016).

We performed downsampling analyses to assess how sequencing depth and the number of cells affected variant classification performance (Methods). This showed that ∼150,000 cells sequenced to a mean depth of ∼5,000 unique molecular identifiers (UMIs) were sufficient to achieve a performance on the ClinVar benchmark and a correlation with PrimateAI-3D scores close to the full dataset (AUROC = 0.97; Figure S4A). Higher numbers of cells improved the correlation between the expression score and PrimateAI-3D predictions independent of sequencing depth (Figure S4B), likely due to better detection of underrepresented or depleted variants, such as TAD I variants that lead to apoptosis and decreased cell proliferation (Figure S4C).

Taken together, these benchmarking results demonstrate the state-of-the-art performance of scRNA-seq as a readout of variant pathogenicity, and enabled classifying 818 out of 2325 missense variants (including 235/559 ClinVar VUSs) as likely pathogenic along the DBD axis and 32 missense variants as likely pathogenic along the TAD I axis on the basis of our assay. 137/818 (17%) of variants with DBD-like pathogenicity were outside of the DBD and 7/32 (22%) of variants with TAD I–like pathogenicity were outside of TAD I, emphasizing the need for a scRNA-seq readout to identify these distinct functional consequences.

### Distinguishing between two axes of pathogenicity in SOD1 improves classification of ALS variants

Despite substantial advances in our understanding of ALS (Mejzini et al., 2019), the precise mechanisms by which SOD1 mutations lead to disease remain incompletely understood. ALS-associated mutations have been identified in almost every region of SOD1 (Figure 5A) (Opie-Martin et al., 2022), with many mutations retaining partial or full dismutase enzyme activity (i.e., converting superoxide to water and hydrogen peroxide). No correlation has been observed between a reduction in dismutase activity, which is essential for cell survival (Tsherniak et al., 2017), and disease age of onset or rate of progression, leading to a consensus that disease likely arises from toxic gain-of-function effects from misfolding or intracellular aggregation that are independent of enzymatic activity (Taylor et al., 2016). Therefore, we sought to identify whether variants that produce these two orthogonal effects could be separated into distinct axes of pathogenicity by their transcriptomic signatures in our single cell assay. We at first used the same experimental design as for p16 and TP53, transducing the SOD1 variant library into an A549 wild type background, but did not detect any variant effects on gene expression or cell proliferation. Reasoning that this was likely due to the absence of dominant negative effects of pathogenic SOD1 variants combined with high endogenous expression, we knocked out endogenous SOD1 using CRISPR prior to lentiviral transduction of the variant library (Methods). The knockout efficiency, assayed using Sanger sequencing of the edited regions, was ∼77-100% across experiments (Methods). We obtained 92,465 cells expressing all 1,395 possible SOD1 SNVs using this approach, with 1,387 variants (99.4%) observed in at least 20 cells, corresponding to 1,106 of 1,113 possible amino acid variants (Figure S5A). Unsupervised two-dimensional projection of variants’ expression profiles showed a gradient of pathogenicity for missense variants that correlated with PrimateAI-3D computational predictions. However, unlike for p16 and TP53, known ALS pathogenic variants in SOD1 did not show a distinct expression signature that was apparent in the unsupervised analysis (Figure S5B).

**Figure 5.**
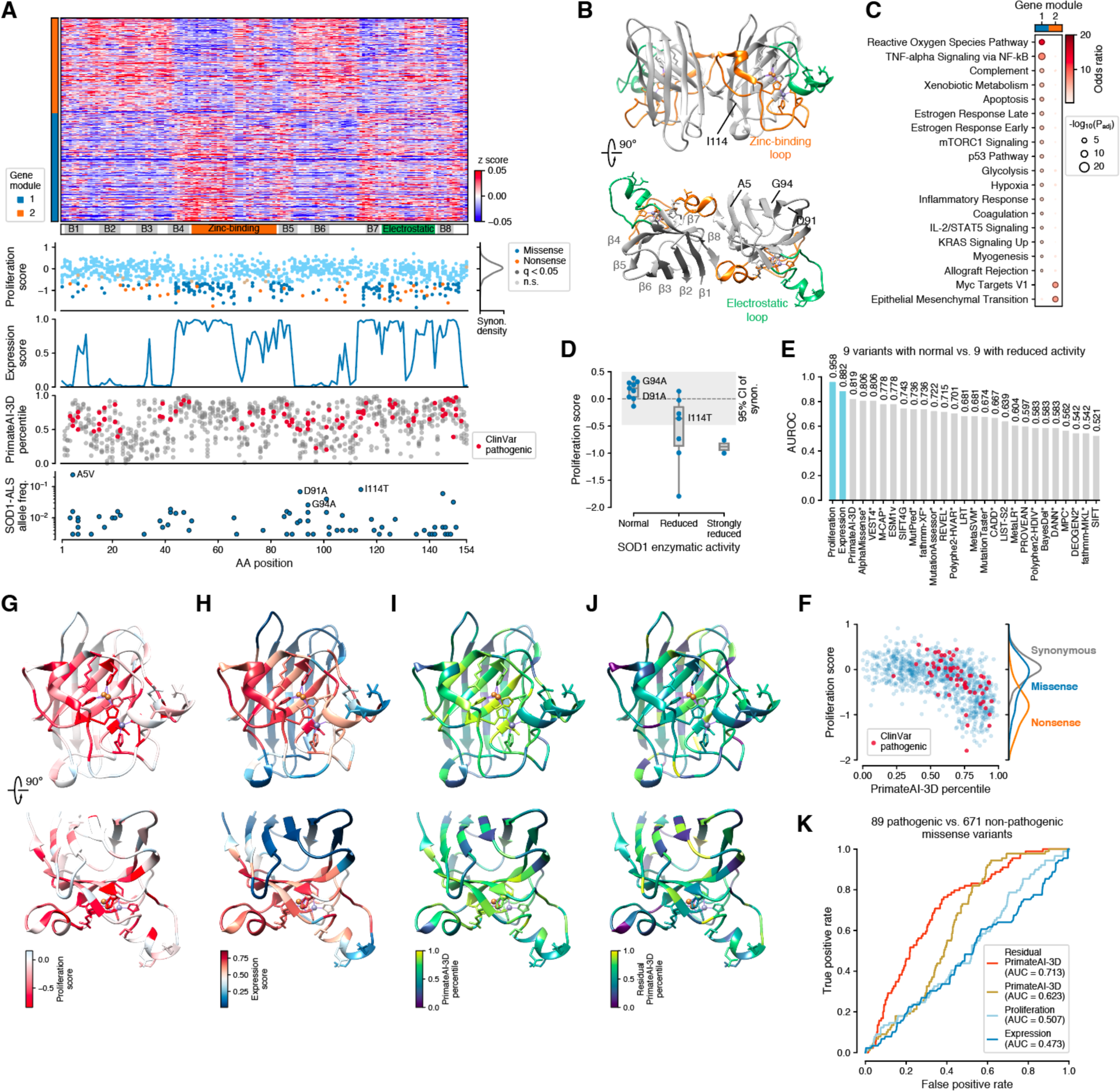
Identification of distinct modes of SOD1 pathogenicity improves classification of ALS variants. (A) Top: heatmap of expression profiles averaged across neighboring residue positions with gene modules from unsupervised clustering (Methods). Bottom panels: proliferation score for missense and nonsense variants, with significantly depleted variants labeled (q-value < 0.05; Methods); expression score; PrimateAI-3D scores for all missense variants, with ClinVar pathogenic variants labeled; frequency of SOD1 variants in a cohort of 1,003 ALS cases (Opie-Martin et al., 2022), with the most common pathogenic variants labeled. (B) Structure of the WT SOD1 homodimer (PDB: 2C9V), with zinc-binding and electrostatic loops and beta barrel labeled. (C) Biological pathways enriched for gene modules up- or down-regulated upon expression of SOD1 variants (Methods). (D) Proliferation scores from our assay compared to previously reported variant effects on enzymatic activity from the literature (Table S4). (E) Performance of single cell assay scores from this study (blue) and computational models (gray) on the task of classifying variant effects on SOD1 enzyme activity, with models that were trained on ClinVar or other clinical variant annotations or human allele frequencies databases indicated by *. (F) Scatter plot of SOD1 missense variants showing the relationship between PrimateAI-3D predictions and proliferation score, with marginal distributions of different types of variants shown for comparison. Note that pathogenic ALS variants (labeled in red) have higher PrimateAI-3D scores than would be expected based on their proliferation score. (G-J) SOD1 monomer (same structure and orientation as in (B)) colored by proliferation score, expression score, PrimateAI-3D score, and residual PrimateAI-3D score after regressing out proliferation and expression scores (Methods). (K) Classification performance for distinguishing ClinVar pathogenic variants from variants not observed in any annotation of likely pathogenic variants (Methods).

We next computed expression profiles for each amino acid position, averaging across neighboring residues in the 3D structure to reduce noise in gene expression (Methods), and trained a classifier to distinguish between variants with synonymous- and nonsynonymous-like expression profiles (Figure 5A, Table S3 and Methods). The resulting expression score revealed that pathogenic variants were predominantly localized to regions of the SOD1 protein that are essential for dismutase function, including the zinc-binding and electrostatic loops and the beta barrel parts adjacent to these domains, specifically the β4, β5, β7, and β8 sheets (Figures 5A and 5B). Variants in these regions were associated with upregulation of reactive oxygen species pathway genes (gene expression module 1, Figures 5A and 5C) and genes in apoptosis-related pathways, underscoring the requirement of SOD1 enzymatic activity for cell viability (Tsherniak et al., 2017). Residues that showed effects on proliferation were clustered in close proximity to the catalytic site (Figure 5G), and cell proliferation was markedly reduced for variants with high expression scores (Spearman’s ρ = -0.59).

Based on the discrepancy with known ALS pathogenic variants in SOD1, together with prior observations that ALS-associated variants typically alter β-barrel folding and dimer contacts rather than catalysis (Bruijn et al., 1998; Deng et al., 1993), we reasoned that the structure-function relationships seen in our experiments likely only measured pathogenicity with respect to enzymatic activity, rather than ALS-associated toxic gain-of-function effects. To confirm this, we curated measurements of SOD1 enzymatic activity for 18 amino acid variants from the literature (Table S4, Methods), and compared these to our proliferation scores (Figure 5D). This showed that most variants with reduced or disrupted enzymatic activity adversely affected cell viability in our assay, with proliferation score distinguishing between variants with reduced vs. normal dismutase activity with high accuracy (ROC AUC = 0.958, Figure 5E). In contrast, most computational models performed poorly at predicting pathogenicity related to reduced dismutase activity (Figure 5E).

Because both types of SOD1 pathogenic mutations, whether they act through enzyme activity or ALS, would be subject to negative selection due to their respective effects on cell viability or early mortality, SOD1 sequence conservation should reflect the sum of both axes of pathogenicity, as should computational predictions, which are closely related to protein sequence conservation. Indeed, we observed a nonlinear relationship between our assay scores and PrimateAI-3D predictions (Figure 5F), suggesting that adjusting the PrimateAI-3D predictions (Figure 5I) by subtracting out the contribution of enzyme-related pathogenicity (Figures 5G and 5H) would create a classifier exclusively focused on the task of predicting ALS-related pathogenic variants. We created residual PrimateAI-3D scores by regressing out the expression and proliferation scores for each variant, which increased pathogenicity predictions for variants in known ALS-associated regions in the β-barrel (Figure 5J) compared with the raw scores (Figure 5I).

We next sought to benchmark these adjusted scores on the task of distinguishing ALS-associated pathogenic variants from benign ones. However, the highly variable penetrance, age of onset, and disease course of ALS observed across SOD1 mutations (Opie-Martin et al., 2022) and absence of common SOD1 missense variants in the population hinder the identification of benign variants, with none annotated in ClinVar. Moreover, SOD1 variants observed in human populations were predominantly very rare and contained a high proportion of pathogenic variants (40%; Figure S5C), hindering their use as putatively benign variants. We therefore benchmarked the scores on the task of distinguishing between ClinVar pathogenic variants and variants that were not annotated as potentially pathogenic in any other database or cohort (Abel et al., 2012; Chen et al., 2021; McCann et al., 2021; Opie-Martin et al., 2022; Ruffo et al., 2022; Yamashita and Ando, 2015). While proliferation alone (i.e., effects on enzymatic activity) was not informative for classifying ALS pathogenic variants (ROC AUC = 0.507), regressing out these effects from PrimateAI-3D and other computational predictions indeed consistently improved ALS variant classification (Figures 5K, S5D and S5E, ROC AUC = 0.713 for the residual score vs. 0.623 for the raw score). 24 out of 27 computational scores achieved higher AUROCs after residualization, including the scores from all 9 methods that did not train on clinical variant annotations (Benjamini-Hochberg adjusted p-value < 0.05, Mann-Whitney U test; Figure S5E).

In summary, our results show that while SOD1 contains two axes of pathogenicity, our assay specifically captured pathogenic effects related to the disruption of SOD1 dismutase function. Although we were not able to directly assay ALS-related effects, due to the contribution of both ALS- and dismutase-associated pathogenicity to computational scores, the scores from our assay nonetheless contributed to improving prediction of ALS-associated pathogenicity.

## Discussion

While all benign variants are alike, each pathogenic variant is pathogenic in its own way. Here, we demonstrate that saturation mutagenesis combined with single cell sequencing can be used as a general-purpose variant interpretation assay that achieves state-of-the-art variant classification performance, while also supplying detailed mechanistic insights through the resulting gene expression signatures. Importantly, our results show that the variant-level transcriptomic readout enables the identification of independent modes of pathogenicity within each gene. In the three genes we studied, our single cell assay revealed a complex diversity of gene expression profiles, including those associated with pathogenic variants affecting protein-protein interactions (p16-CDK6 and TP53-MDM2), DNA-binding (TP53-DBD), apoptosis (TP53-TAD I), tetramerization (TP53-4D), enzymatic activity (SOD1), and toxic protein aggregation (SOD1). In the example of SOD1, we were also able to show that isolating the independent contributions of ALS and enzymatic variants improves computational predictions of ALS variants. At the same time, we observe that the different types of pathogenicity present in a gene can confound computational models, particularly when relying on sequence conservation, which only shows that variants at a certain position in the protein are likely to be rejected by natural selection, without being able to distinguish the underlying causes.

Although our results demonstrate the power of using single-cell transcriptomes to characterize variant pathogenicity, our study also identified important limitations with this approach. First, as our initial results with SOD1 suggest, simple overexpression of variant transcripts is likely to work only for variants and genes with dominant negative effects or low endogenous expression. Performing a knockout of the endogenous copies prior to transduction of the variant library is a feasible solution but poses additional challenges that may further complicate the experimental design. Several alternative approaches exist for introducing single variants into a locus, including saturation genome editing, which has been used in proliferation-based deep mutational scanning experiments (Findlay et al., 2018), base editing (Bock et al., 2022) and landing pad systems (Matreyek et al., 2017), and while these each have their limitations at present, it may be feasible to adapt them to perform single cell profiling at greater scale. Second, while we showed that random variant barcoding combined with amplicon sequencing yields highly accurate variant calls, short reads currently limit the assay to the ∼65% of genes that are at most ∼1300-1500 nucleotides long. As long read technologies become more scalable this limitation will disappear; alternatively dual barcoding at the 3’ and 5’ UTRs should enable characterizing up to 90% of all protein coding genes. Third, as our results for SOD1 showed, it may not be possible to detect disease-relevant phenotypes or pathogenic effects in the transcriptome of arbitrary cell lines or at the time scales (days to weeks) that saturation mutagenesis experiments are typically conducted in. However, these limitations are independent of the specific type of assay and readout used, and selecting the right conditions is beyond the scope of this work. Lastly, due to the high level of noise and sparsity in single cell gene expression profiles, hundreds of thousands of cells must be profiled to distinguish variant- and domain-specific pathogenic effects. While the high costs of droplet-based single cell library preparation methods are currently a limiting factor, split-pool approaches that enable characterizing hundreds of thousands of cells in a single experiment (Bock et al., 2022), combined with rapidly diminishing costs of sequencing, will enable scaling these experiments to hundreds of genes. Despite these limitations, the functional interpretation enabled by the scRNA-seq readout of our assay overcomes the major limitation of current experimental assays and computational models, which, by relying on measures of constraint only, inherently reduce the characterization of pathogenicity to a single dimension.

In addition to the advantages described above, single cell profiling also has the potential to streamline mutation scanning experiments by eliminating the requirement of designing bespoke assays for each individual gene. Hence, we anticipate that as the costs of sequencing fall, single cell profiling will increasingly become the preferred readout for saturation mutagenesis experiments. Profiling large numbers of genes, at least enough to cover representatives of all major human protein domains, will in turn drive training of computational models that can predict different modes of pathogenicity together with their effects on the transcriptome. Shifting towards multi-dimensional characterization of variant pathogenicity will enable our field to move beyond the limitations of assessing variant effects only along a single dimension within each gene, substantially enriching our understanding of gene function, molecular pathways, and the consequences of genetic perturbations in health and disease.

## Supplemental Figures

**Figure S1.**
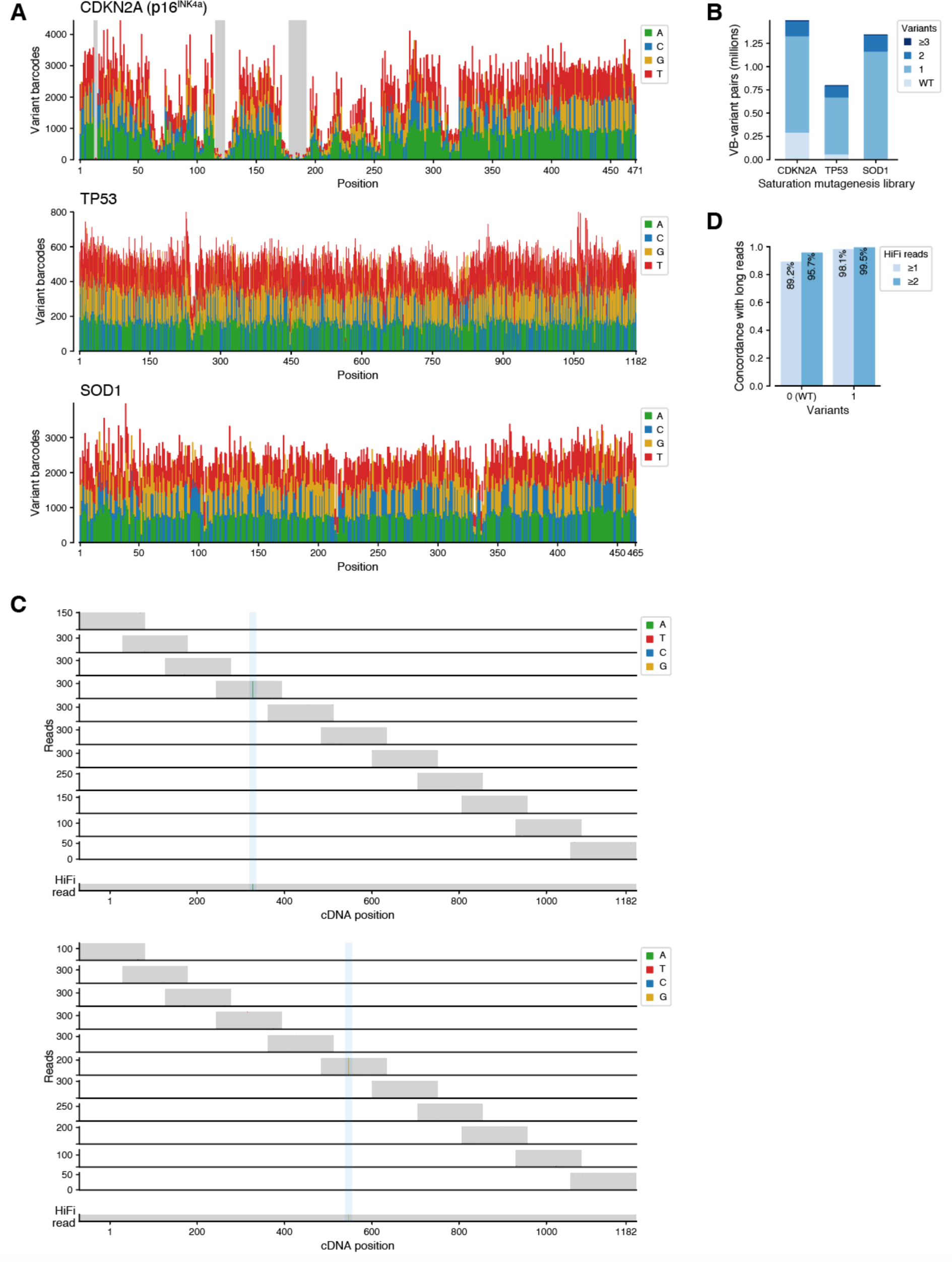
Saturation mutagenesis libraries and validation of amplicon sequencing with long reads. (A) Coverage of each library from amplicon sequencing, shown as number of variant barcodes per alternative base at each position. Shaded areas for CDKN2A indicate positions with poor coverage that were excluded from analyses. (B) Number of variant barcode–variant combinations in each library as detected by amplicon sequencing. Each library contained variant barcodes associated with the wild type (WT) sequence or multiple variants in addition to single variants. (C) Examples of amplicon sequencing data for two TP53 variants, with pileups from each amplicon shown in the top panels and the corresponding HiFi long read (PacBio) in the bottom panel. (D) Concordance of variant calls from amplicon sequencing with HiFi long reads, for variant barcodes with the WT sequence or 1 variant.

**Figure S2.**
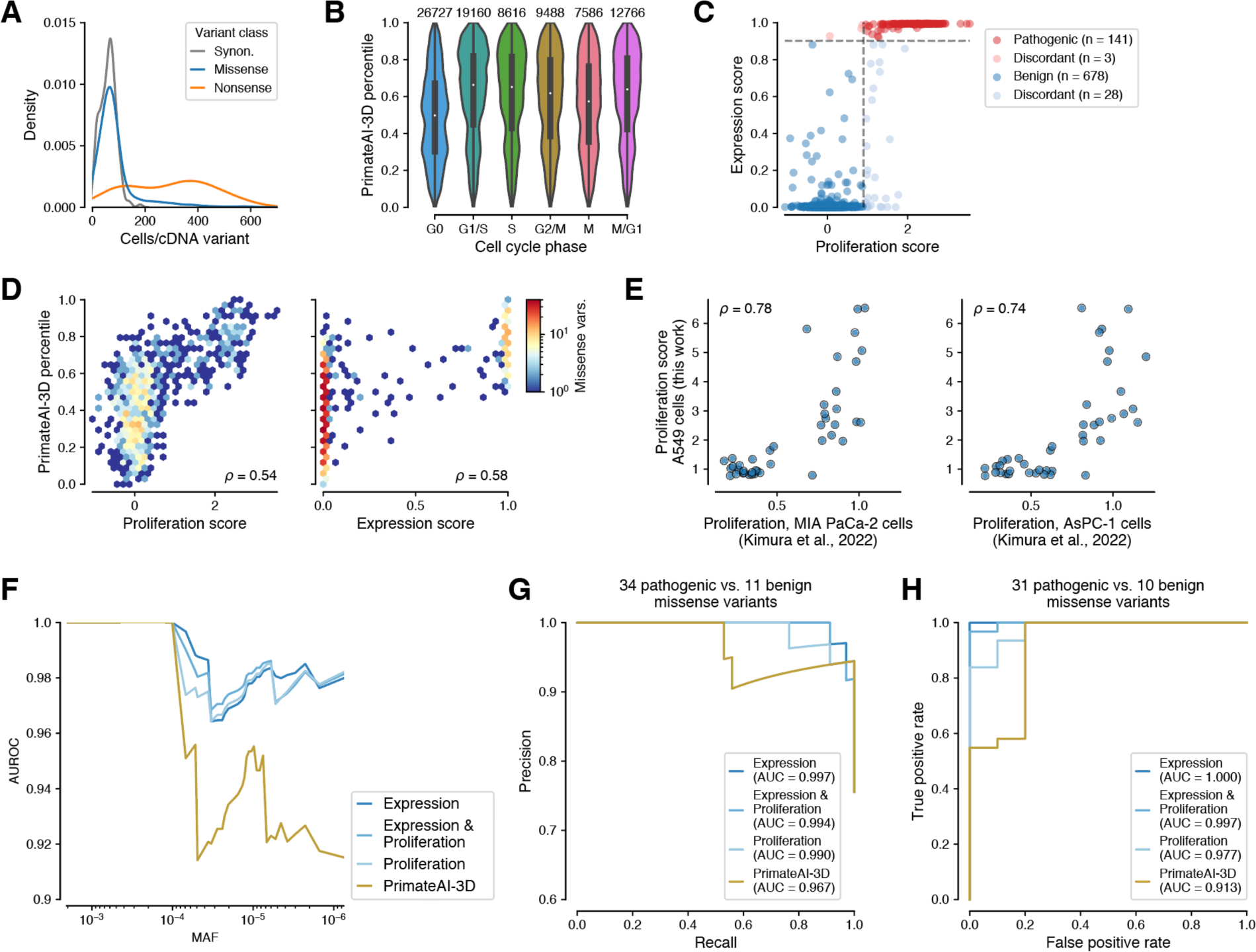
Properties of p16 pathogenicity scores from saturation mutagenesis in single cells. (A) Distribution of the number of cells obtained per cDNA variant for synonymous (synon.), missense, and nonsense variants. (B) Distribution of PrimateAI-3D scores stratified by cell cycle phase, with cell numbers indicated at the top. (C) Concordance between the expression and proliferation scores, with dashed lines indicating the 0.05 FDR threshold for each score for classifying pathogenic variants. (D) Correlation of the proliferation and expression scores with pathogenicity predictions for missense variants from PrimateAI-3D. (E) Correlation of the proliferation score from A549 lung carcinoma epithelial cells (this work) with proliferation scores from two pancreatic ductal adenocarcinoma cell lines for 43 variants assayed in (Kimura et al., 2022). (F) Performance of experimental scores and computational predictions on the task of distinguishing 31 pathogenic variants in ClinVar from 4 benign variants supplemented with variants observed in human populations up to a selected minor allele frequency (MAF) threshold. (G) Precision-recall curves and area under the curve (AUC) for single cell gene expression and proliferation-derived scores and computational predictions for classifying ClinVar pathogenic from benign variants. (H) ROC curves and AUC for single cell gene expression and proliferation-derived scores and computational predictions for distinguishing ClinVar pathogenic from benign variants, for variants observed in at least 30 cells.

**Figure S3.**
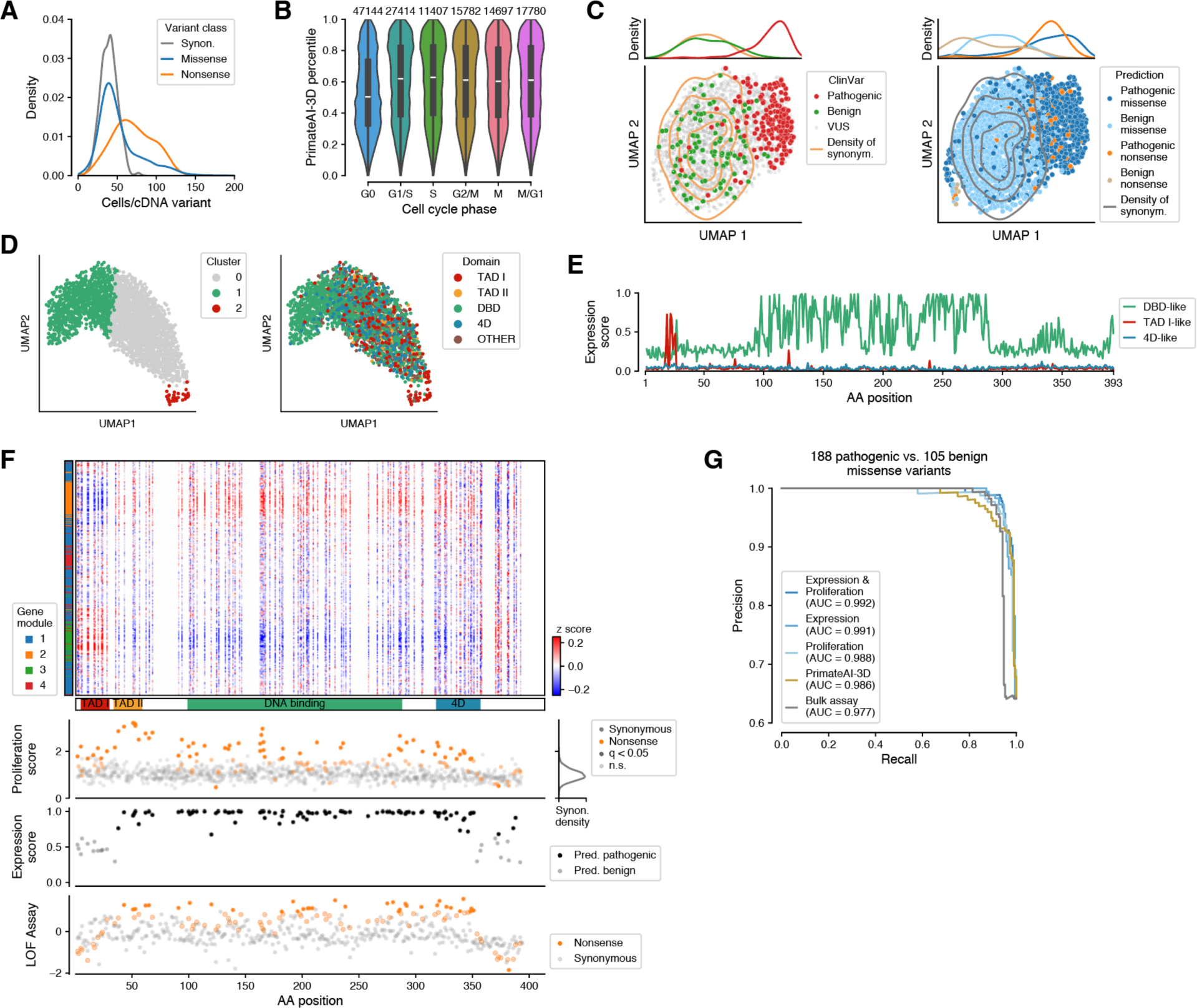
Properties of domain-specific modes of pathogenicity in TP53. (A) Distribution of the number of cells obtained per cDNA variant for synonymous (synon.), missense, and nonsense variants. (B) Distribution of PrimateAI-3D scores stratified by cell cycle phase, with cell numbers indicated at the top. (C) UMAP of expression profiles averaged by amino acid, labeled by ClinVar annotation (left) and by the predicted pathogenicity from the ensemble classifier (right), with the distribution of synonymous variants shown as density contours and the densities along UMAP dimension 1 shown at the top. (D) UMAP of expression profiles averaged by residue position, labeled by unsupervised clusters (left) and domain annotations (right). (E) Expression scores for variants with an expression signature matching that of variants in the DBD, in TAD I, or in 4D, showing that no variants had a signature specific to the tetramerization domain. (F) Top: heatmap showing the expression profile (z-score) of nonsense variants averaged across neighboring residue positions (Methods). Bottom panels: proliferation score for nonsense variants, with significantly enriched or depleted variants indicated (q-value < 0.05); expression score for nonsense variants, with variants predicted to be pathogenic indicted (q-value < 0.05); loss-of-function bulk assay score from (Giacomelli et al., 2018). (G) Precision-recall curves and area under the curve (AUC) for the benchmark from Figure 4A.

**Figure S4.**
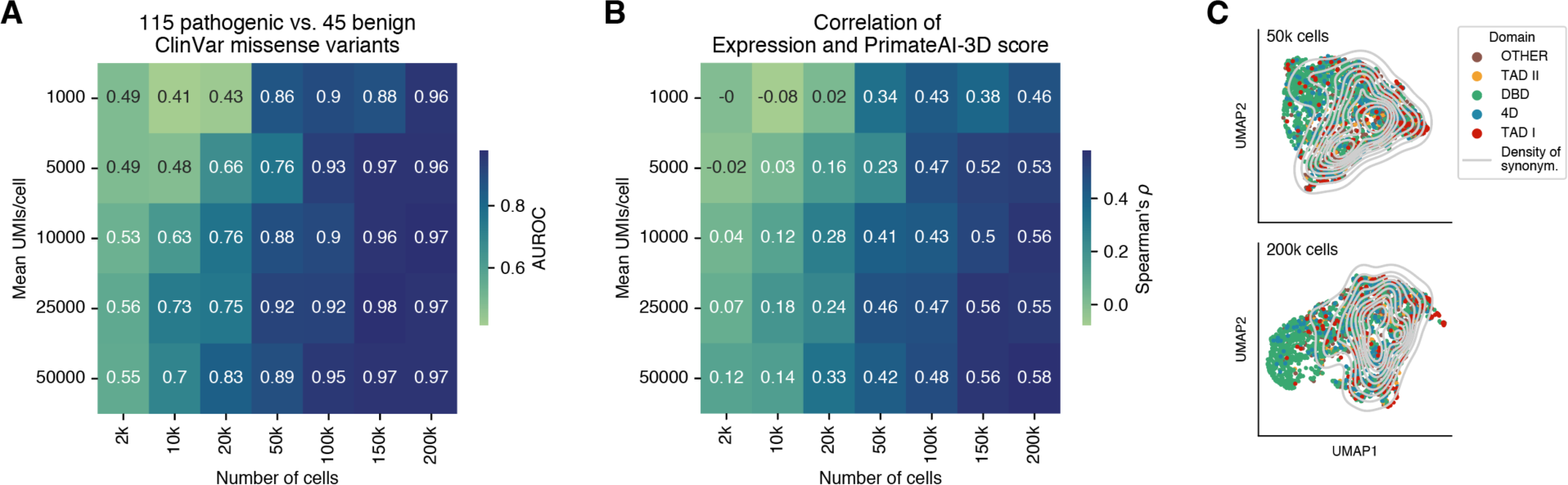
Variant classification performance as a function of sequencing depth and number of cells. (A) Classification performance (AUROC) on the task of distinguishing pathogenic from benign ClinVar variants as a function of total cell number and mean sequencing depth per cell, for the subset of variants detected across all downsampling conditions. (B) Spearman correlation between expression score (i.e., the probability of a variant having DBD- or TAD I–like pathogenicity based on its expression profile) and predicted pathogenicity from PrimateAI-3D. (C) UMAP of amino acid–level average expression profiles computed from a total of 50k (top) and 200k cells (bottom) sampled to a mean of 50k UMIs, showing that larger cell numbers improve the detection of DBD- and TAD I–like pathogenic effects. UMI: unique molecular identifier.

**Figure S5.**
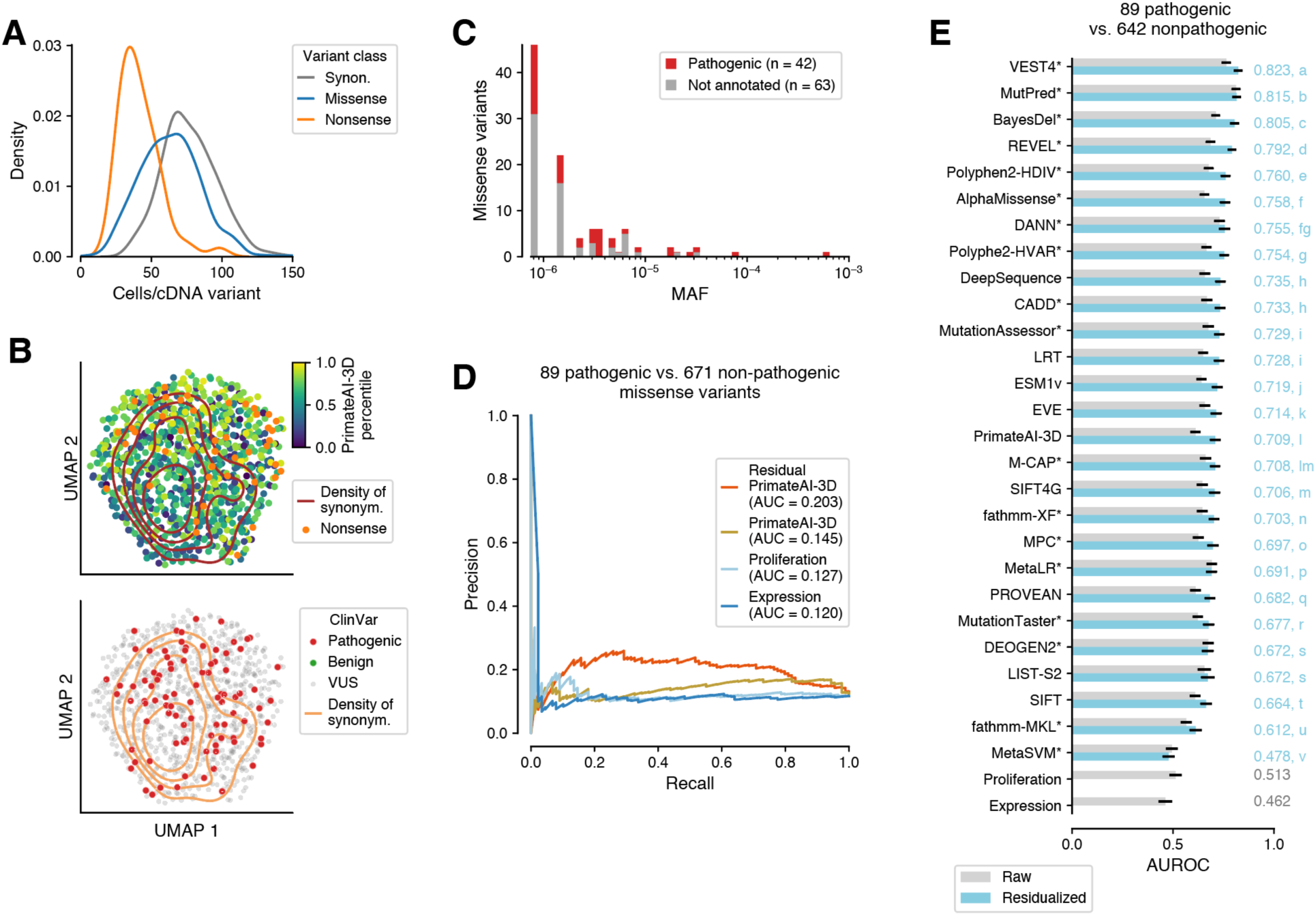
Benchmarking of catalysis- and ALS-associated pathogenicity in SOD1. (A) Distribution of the number of cells obtained per cDNA variant for synonymous (synon.), missense, and nonsense variants. (B) UMAP of expression profiles averaged by amino acid for missense variants labeled with PrimateAI-3D scores, with the distribution of synonymous variants shown as density contours (top) and labeled with ClinVar annotations (bottom). (C) Allele frequency of SOD1 missense observed across ∼416,800 individuals in UK Biobank, gnomAD, and TOPMed, containing 42 of the 56 pathogenic variants annotated in ClinVar. (D) Precision-recall curves and area under the curve (AUC) for the benchmark from Figure 5J. (E) Performance of experimentally derived scores and computational models on classifying pathogenic vs. benign ClinVar variants, for raw scores and scores residualized by the expression and proliferation scores.

## Methods

### Key resources table

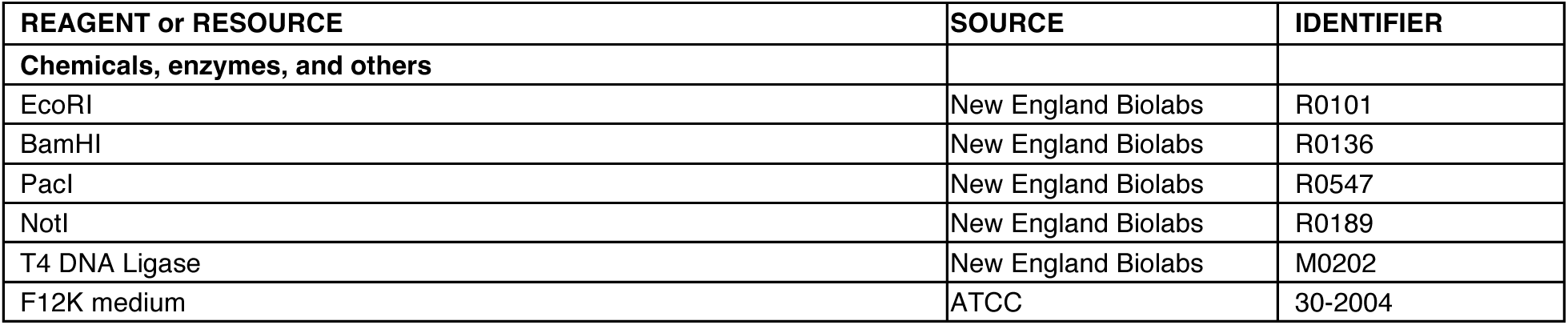
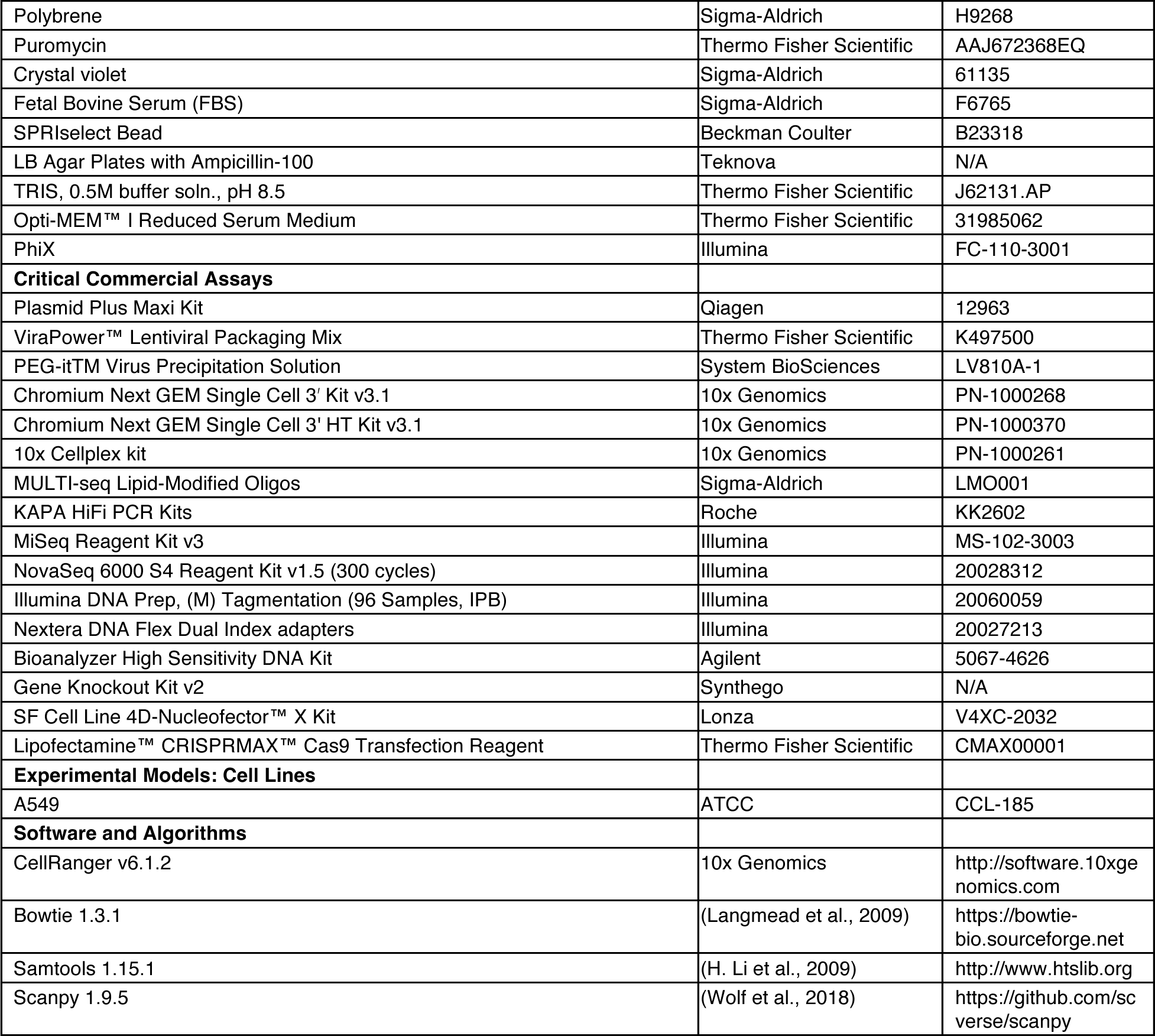

### Construction of variant libraries

To generate variant libraries, plasmids containing wild type ORFs were obtained from GenScript. The ORFs were cloned into the pcDNA3.1(-) backbone (Thermo Fisher Scientific V79520) between the EcoR1 and BamH1 sites, with the addition of a Kozak sequence (GCCACC) before the start codon. The wild type sequences used were NM000546 for TP53, NM00007 for CDKN2A, and NM000454 for SOD1. The constructed plasmids were then submitted to Twist Bioscience for the subsequent synthesis of the variant libraries. Flanking sequences (upstream: gccctctagactcgagcggccgccactgtgctggatatctgcagaattcgccacc, downstream: ggatccgagctcggtaccaagcttaagtttaaacc) were incorporated from the pcDNA3.1(-) vector to serve as cloning handles. The libraries for TP53 and SOD1 contained all 3,546 and 1,395 possible SNVs, respectively.. However, the CDKN2A library failed to pass Twist Bioscience’s internal quality controls for variants at codon 5, 41, 61, and 63. Due to low coverage of variants around these positions (Figure S1A), we excluded variants at positions 5, 39-41, and 60-64.

### Construction of 3’ UTR barcoded lentiviral expression vector

The lentiviral expression vector sequence from (Ursu et al., 2022) was modified by deleting the original Not1 and BamH1 sites and adding a Multiple Cloning Site (MCS) for library cloning. This lentiviral plasmid was synthesized by GenScript. To create the barcoded lentiviral vector, a 19 bp semi-random barcode (BNSWNSKMSVNBSWNMKYN) was inserted into the MCS between the BamH1 and PacI sites using Gibson assembly. The semi-random sequence was designed to minimize homopolymer formation and AATAAA generation. The barcoded vector was transformed into E. coli and subsequently plated on agar plates. After overnight incubation, bacterial DNA from around 4 million colonies was then extracted to achieve high barcode diversity. To assess barcode diversity, we performed paired-end sequencing of a small amplicon covering the barcode region, resulting in over 3.8 million different variant barcodes.

### Cloning of variant libraries into barcoded lentiviral vector

The variant libraries of CDKN2A, TP53, and SOD1 were cloned into the barcoded lentiviral vector by restriction digestion and ligation between the Not1 and BamH1 sites on the MCS. The cloned libraries were transformed into E. coli and subsequently plated on agar plates. After overnight incubation, library DNA from over 4 million different colonies was extracted to ensure high barcode diversity and full representation of the variants, with amplicon sequencing as the quality control method. All variants synthesized by Twist Bioscience were successfully cloned into the vector, resulting in over 750,000 different variant barcode-variant combinations for each gene.

### Lentiviral production and titering

Lentivirus production was carried out using the ViraPower™ Lentiviral Packaging Mix (ThermoFisher Scientific K497500) in the 293FT cell line (ThermoFisher Scientific R70007) following the manufacturer’s protocol. The collected virus was concentrated 100-fold after precipitation with PEG-itTM Virus Precipitation Solution (System BioSciences LV810A-1) for 3 days. The titer of the lentiviral library was determined by transducing A549 cells (ATCC CCL-185) with the lentivirus at different dilutions in 1-ml F12k media (ATCC 30-2004) in the presence of 6 ug/ml polybrene (Sigma H9268). After selection with 1.5 ug/ml puromycin (ThermoFisher AAJ672368EQ) for 10 days, the number of puromycin-resistant colonies was counted using crystal violet (Sigma 61135) staining. The viral titer used in this study was determined to be 1.35*107 IFU per ml for TP53, 1.88*107 IFU per ml for SOD1, and 1.29*107 IFU per ml for CDKN2A.

### Lentiviral transduction with variant libraries

A549 cells cultured in F12k medium with 10% FBS (Sigma) were seeded at around 10% confluency on day 1. On day 2, the packaged lentiviral stocks containing the variant libraries were added to the cells at a MOI ranging from 0.003 to 0.1, along with 6 ug/ml of polybrene (Sigma-Aldrich, H9268). The medium was replaced on day 3 to remove the polybrene. On day 4, 2 ug/ml of puromycin was added to the medium to select the successfully transduced cells. After 3 days of puromycin selection, the cells were recovered in medium without puromycin for 2-3 days before being harvested for 10x Genomics single-cell RNA-seq library preparation.

### Single-cell RNA-seq library preparation and sequencing

Standard loading single-cell RNA-seq libraries were prepared using the 10x Genomics Chromium Next GEM Single Cell 3ʹ Kit v3.1 (PN-1000268) or the Chromium Next GEM Single Cell 3’ HT Kit v3.1 (PN-1000370) on the Chromium controller or Chromium X. For some libraries, the 10x Cellplex kit (PN-1000261) or MULTI-seq workflow (McGinnis et al., 2019) was used for barcoding and overloading, allowing for the identification of multiplets during GEM formation. The final libraries were sequenced on NovaSeq 6000 systems, generating a minimum of 40,000 reads on average per cell. The sequencing length was 28 cycles for read 1 and 150 cycles for read 2.

### Amplicon sequencing to link variants to 3’ UTR barcodes

cDNA samples were generated during the 10x library preparation, 10 ng of which were used as a template for a two-step PCR to connect variants with variant barcodes. In the first PCR, each reaction uses a reverse primer mixture consisting of an equal volume of four staggered oligos that annealed to the vector backbone in the 3’ region of the barcode (i.e., 3BC B15 0N, 3BC B15 1N, 3BC B15 2N, 3BC B15 3N). To cover the 1182, 471 and 465 nucleotide long sequences of TP53, CDKN2A and SOD1, respectively, with tiling 150 bp reads, 11 amplicons were required for TP53 and 4 amplicons for CDKN2A and SOD1. For each amplicon, a gene-specific forward primer was added separately to each PCR reaction (the specific sequences are provided below). The final concentration of primers in all PCRs was 0.25 uM. The PCR products were purified using 0.8x SPRIselect beads and eluted in 52.5 ul of 10mM Tris buffer solution, pH 8.5. In the second PCR, which is an indexing PCR, 5 ul of the purified products from the first PCR were used as the template, along with 10 ul of Nextera DNA Flex Dual Index adapters (Illumina: IDT for Illumina DNA/RNA UD indexes: 20027213) as primers. The PCR products were purified using 1x SPRIselect beads and eluted in 27.5 ul of 10mM Tris buffer solution, pH 8.5. Both PCRs utilized 2x KAPA HiFi HotStart ReadyMix (Kapa KK2602). The first gene-specific PCR consisted of 16 cycles (95°C for 3 min, followed by 16 cycles of 95°C for 30 seconds, 60°C for 30 seconds, 72°C for 30 seconds, and a final extension at 72°C for 5 min), while the second indexing PCR consisted of 9 cycles using the same cycling conditions. The eluted PCR products were quantitated using a Bioanalyzer or Tapestation (Agilent). The amplicons were pooled for sequencing on NovaSeq S4 flow cells using 80PM as the loading concentration with the NovaSeq Xp workflow. To increase the diversity of the amplicons, the libraries were mixed with 30% PhiX control libraries (Illumina FC-110-3001) and sequenced for 150 cycles for Read 1 and 100 cycles for Read 2. A minimum of 20 million reads were generated for each amplicon.

The reverse PCR primers on the barcode side consisted of a mixture of staggered primers, with the following sequences used for all genes:

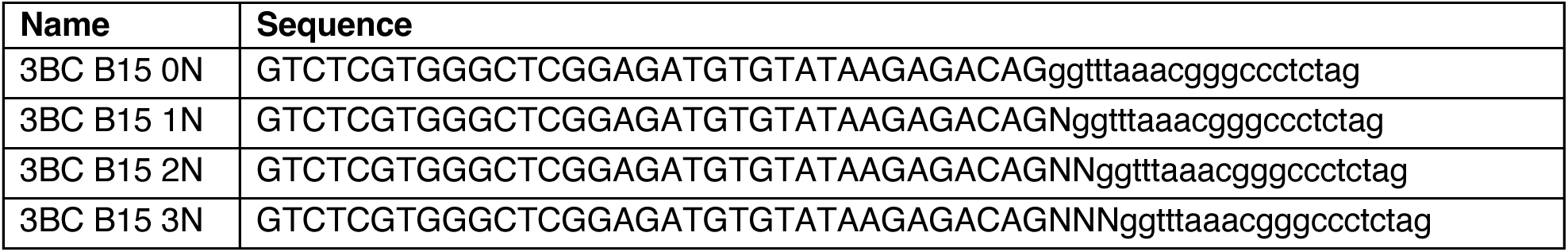

The forward primers on the variant side had the following sequences:

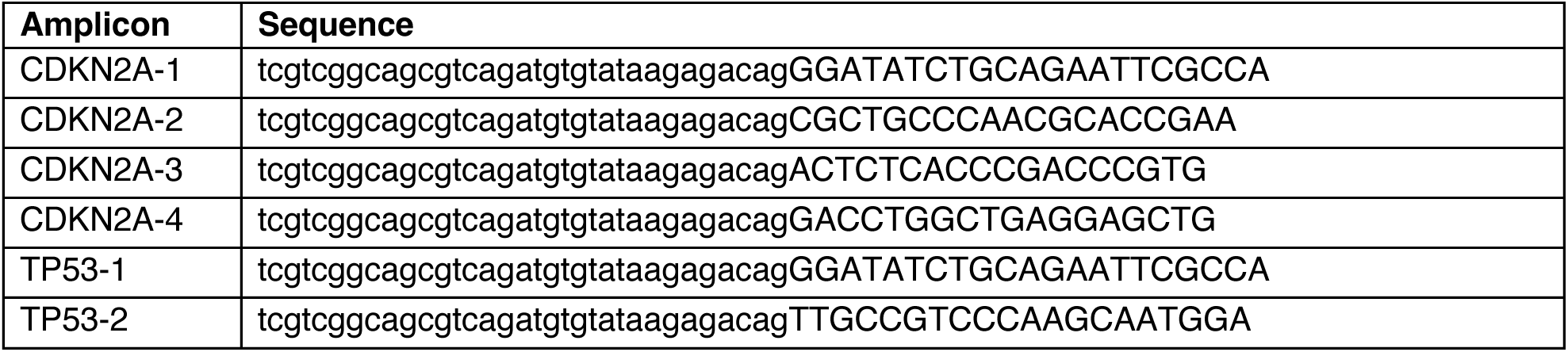
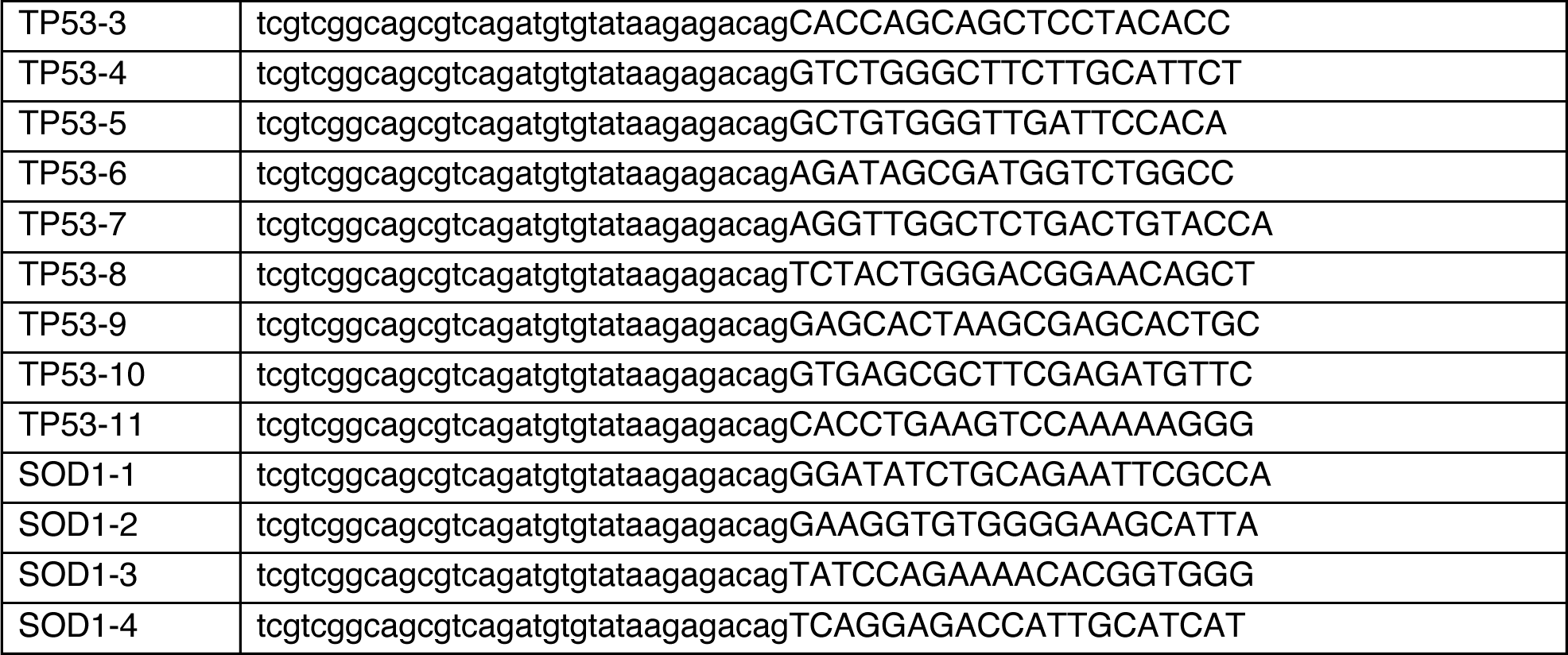

### CRISPR knockout experiments

The SOD1 CRISPR knockout experiments utilized Synthego’s Gene Knockout Kit v2 following the manufacturer’s protocols. Predesigned sgRNAs specific to SOD1 and SpCas9 2NLS nuclease were ordered from Synthego. A549 cells were seeded in culture dishes or plates and incubated to obtain 50-70% confluency at the time of transfection, using either electroporation or chemical transfection. For electroporation, the Lonza 4D-Nucleofector^TM^ System with X unit and 4D-Nucleofector^TM^ X Kit (32 RCT) suitable for A549 cells were used. Briefly, the harvested cells were washed and resuspended in the provided nucleofector solution. Ribonucleoprotein (RNP) was formed using the sgRNA and SpCas9 nuclease, then mixed with the cell suspension. The mixture was then electroporated using the specific electroporation parameters designated by the Lonza system. In parallel, chemical transfection was performed using Lipofectamine^TM^ CRISPRMAX^TM^. As in the electroporation method, the harvested cells were subjected to a washing step before being resuspended. RNP-transfection solution prepared from Opti-MEM medium, Lipofectamine^TM^ CRISPRMAX^TM^ reagent, and the formed RNP was added to the cells and incubated for 3 days to allow for transfection and recovery. The efficiency of SOD1 gene knockout was evaluated by PCR-amplifying the edited region and Sanger sequencing. The .ab1 files from Sanger sequencing from the knockout and control groups were submitted to Synthego’s ICE analysis tool to calculate a knockout score, which represents the proportion of cells that have either a frameshift-inducing indel or a >21bp indel in the protein-coding region and are likely to induce complete loss of function. The knockout scores for the SOD1 experiments ranged from 77-100%.

### Variant barcode mapping from 3’ scRNA-seq libraries

To assign variant barcodes to cells, the FASTQ files generated by CellRanger (mkfastq subprogram) were parsed for reads containing the nucleotide sequence upstream of the variant barcode. These reads were then filtered to ensure that the variant barcode sequence matched the template BNSWNSKMSVNBSWNMKYN as well as the downstream sequence, allowing for up to two mismatches. For each resulting cell barcode-UMI pair, the corresponding reads were checked for concordance with a single variant barcode, allowing up to two mismatches. UMIs with multiple reads and discordant variant barcodes (i.e., likely chimeric molecules created during library preparation) were discarded (typically < 0.1% of UMIs).

### Variant calling from amplicon sequencing

We developed a custom pipeline to call variants from amplicon sequencing. Each amplicon was first processed individually, starting from FASTQs, with read 1 containing the targeted portion of coding sequence and read 2 containing the variant barcode. The variant barcode was parsed from read 2 by determining the offset of the staggered primer sequence and checking the variant barcode and downstream sequences for concordance with the reference template, allowing up to one mismatch. Next, we appended the variant barcode to the read ID in read 1, and aligned this new FASTQ to the wild type reference sequence using Bowtie 1.3.1. From these alignments, we generated pileups of all reads corresponding to a variant barcode, and computed alternate allele frequencies and counts for variants found in at least 3 reads with base quality ≥ 37. Next, we aggregated variant calls across all amplicons, retaining variant barcodes only if they were observed in all amplicons, and if they contained at least one variant (the saturation mutagenesis libraries contained between ∼1-22% of wild type sequences). Variant calls were required to match in regions of overlap between amplicons. Finally, we filtered variants on allele frequency (≥0.6) and total coverage (≥5 reads) to minimize potential template switching artifacts that may have arisen during PCR.

### Validation of amplicon sequencing with long reads

To test the accuracy of amplicon sequencing, we sequenced the TP53 library using PacBio HiFi long reads, produced using circular consensus sequencing. We linearized the TP53 variant library, which included the random variant barcode and mutagenized coding sequences, using Not I digestion of the plasmid DNA, and submitted this for sequencing on the PacBio Sequel® platform to Azenta Life Sciences. We only used reads that contained the full coding and barcode sequences, excluding reads with any insertions or deletions. We obtained HiFi reads matching 135,589 variant barcodes for which we observed a single variant (94.8%) or the WT sequence (5.2%) with amplicon sequencing. Of these, 132,382 (97.6%) matched the amplicon sequencing data. After filtering for the 64,206 variant barcodes that were observed in at least two HiFi reads, the concordance increased to 99.3% (63,773 matching reads).

### Single cell RNA-seq expression quantification and quality control

All scRNA-seq experiments were processed using CellRanger 6.1.2 (10x Genomics) to generate gene expression matrices for downstream analyses. We processed the UMI count matrices generated by CellRanger with Scanpy (Wolf et al., 2018), first normalizing counts in each cell to 10,000, followed by log1p-transformation (i.e., generating log(CP10k+1) units). We annotated the cell cycle phase of each cell as described below. Next, we selected genes with variable expression across cells using scanpy.pp.highly_variable_genes, resulting in 1,690 genes for CDKN2A and 1,797 genes for TP53. For SOD1 we selected the top 5,000 variable genes using the ’seurat_v3’ option. We regressed out the total read count and experiment from the expression matrix of variable genes, treating each experiment as a one-hot encoded batch covariate. Due to the roles of p16 and p53 in regulating cell cycle progression, we did not include cell cycle as part of the covariate correction applied during preprocessing of the single cell gene expression data. Correcting for cell cycle did not modify the results for SOD1, and we therefore also omitted cell cycle from the covariates for SOD1. After covariate correction, gene expression was transformed to z-scores. All analyses were performed using the resulting normalized gene expression matrices with highly variable genes, unless specified otherwise.

To exclude cells with outlier gene expression driven by quality issues rather than variant effects, we performed Leiden clustering with a resolution of 1, based on a neighborhood graph with 15 neighbors. This overclustered the cells, enabling the identification and exclusion of a small number of outlier clusters that had low numbers of UMIs/cell, low numbers of cells, a high percentage of mitochondrial reads (>20% on average), or only represented cells from a minority of experiments.

### Cell cycle annotation

To assign each cell to a cell cycle phase, we adopted the same strategy as (Ursu et al., 2022), based on marker genes for G1/S, S, G2/M, M, and M/G1 from (Macosko et al., 2015). Briefly, we computed the average expression of marker genes (in log(CP10k+1)) for each phase in each cell, filtering out genes with low correlation (Pearson r < 0.3) to the mean, followed by standardization. The phase with the highest score was assigned to each cell; cells with all scores < 0 were assigned to G0.

### Gene module definitions and gene set enrichment analyses

Gene set enrichment analyses were performed by first obtaining unsupervised clusters of genes based on the residue position–level average expression profiles. We used Leiden clustering with 5 nearest neighbors and the resolution selected for each gene to yield clusters or ‘gene modules’ with interpretable differences in the pathway enrichment results (0.2 for CDKN2A, 0.4 for TP53, and 0.1 for SOD1). For each module, we computed gene set enrichments with Enrichr (Kuleshov et al., 2016) and GSEApy (Fang et al., 2023), using the MSigDB_Hallmark_2020 gene set and retaining enrichments with adjusted p-value < 0.05 for further analysis.

### Proliferation scores

Proliferation scores for missense and nonsense variants were computed as the ratio of cells containing a given variant over its abundance in the library, normalized using synonymous variants. This was performed by aggregating the number of cells for each variant across all available experiments. The cell counts were then scaled by the median number of barcodes for synonymous variants in the library, divided by the median number of cells per synonymous variant. To identify variants with a statistically significant change in proliferation, we first computed an empirical p-value for each missense or nonsense variant based on the distribution of proliferation scores for synonymous variants. We then used these p-values to compute q-values (Storey and Tibshirani, 2003) with the ‘lambda’ parameter set to 0.5 (i.e., using the p-value interval [0.5, 1] to estimate the proportion of true null p-values, ρχ_0_), and applied a 0.05 FDR threshold.

### Expression-derived pathogenicity scores

To generate pathogenicity scores from expression, we performed unsupervised clustering and trained supervised models to predict cluster labels.

To generate the cluster labels for CDKN2A, we excluded amino acid variants with insufficient library coverage and low cell counts (<30 cells) and then performed UMAP embedding on missense and synonymous variants using the first 50 PCs of the amino acid–level average gene expression profiles computed using all available cells for each variant. We used 20 nearest neighbors and a minimum distance between embedded points of 1. We then applied Leiden clustering with a resolution of 0.2 on the UMAP neighborhood graph, resulting in two distinct clusters of variants, which we labeled as ‘Synonymous’ and ‘Non-synonymous’, respectively, given that the former was grouped with synonymous variants. For TP53, we calculated average expression profiles for each amino acid variant using all available cells. These profiles were then averaged by position and further averaged within a 12Å radius in the TP53 AlphaFold structure. We then performed UMAP embedding as described for CDKN2A, but with a minimum distance of 1.5, followed by Leiden clustering with a resolution of 0.2 to identify variant clusters. Clusters were labeled using majority voting; if a cluster predominantly contained synonymous variants, all variants within it were labeled as ‘synonymous’, and if the majority represented the DBD, the cluster was labeled as ‘DBD’. This process resulted in four distinct classes: Synonymous, TAD I, DBD, and 4D variants. For SOD1, we followed the same procedure but limited averaging across neighboring residues in the SOD1 AlphaFold structure to a radius of 6 Å. We performed UMAP embedding as described for CDKN2A, followed by Leiden clustering with a resolution of 0.25, resulting in three clusters of variants. We labelled the cluster overlapping synonymous variants as ‘synonymous’, and the other two as ‘non-synonymous’.

Next, we trained XGBoost models separately for each gene with hyperparameter selection to predict variant classes. We used averaged expression of amino acid variants from all available cells for CDKN2A and TP53 as input, and augmented the synonymous class with wild-type samples (averaged across randomly selected, non-overlapping samples of 200 cells). For SOD1, due to the higher level of noise, we used the expression averaged across neighboring residues as the input. We held out 10% of the data for testing. For the remaining 90%, we trained an XGBoost model with hyperparameter selection with five-fold cross validation using multi:softprob as the objective. To utilize the gradient in the expression profile, we added sample weighting when training the TP53 model. Sample weights for missense variants were calculated based on their distance from the center of the first two principal components of synonymous variants divided by the median distance of all missense variants from the synonymous center, ensuring higher weights for variants farther away from synonymous variants. WT and synonymous samples were assigned a sample weight of 1.

We selected the best model based on minimal multi-class log loss of the cross-validation runs. Predictions for the held-out test set, nonsense variants, and missense variants not included in training (CDKN2A variants with fewer than 30 cells) were averaged across five models from the five folds. We refer to the predicted probability of being in a class as the expression score for that class. For CDKN2A and SOD1, the expression score is the probability of being non-synonymous. For TP53, in addition to expression scores for TAD I, DBD, and 4D, a general expression score was defined as 1 - probability of being synonymous.

We computed the ensemble score (Expression & Proliferation) by normalizing the proliferation score to the unit interval, adding it to the expression score, and normalizing the resulting sum to the unit interval. Since TP53 TAD I variants were associated with reduced proliferation, we instead started by normalizing the negative proliferation score to the unit interval for these variants.

To select variants with expression scores or the ensemble scores that differed significantly from synonymous variants, we calculated an empirical p-value for each variant based on the distribution of scores for synonymous variants, computed adjusted p-values using Benjamini-Hochberg to correct for multiple testing, and classified variants with adjusted p-values < 0.05 as pathogenic.

### Variant annotations

We used HGVS conventions (den Dunnen et al., 2016) for all variant descriptions, omitting the ‘p’ prefix for amino acid variants for simplicity.

### Benchmarking

ClinVar annotations for the three genes were downloaded from https://www.ncbi.nlm.nih.gov/clinvar/ on 09/30/2023 and filtered for coding SNVs, excluding VUSs. We also excluded variants with SpliceAI scores > 0.5 (Jaganathan et al., 2019), for example the c.457G>T mutation in CDKN2A (Rutter et al., 2003), since pathogenic effects from alternative splicing were not captured in our experiments. For all benchmarks, we merged variants annotated as ‘Pathogenic’, ‘Pathogenic/Likely pathogenic’ or ‘Likely pathogenic’ into a single ‘Pathogenic’ category, and similarly, variants annotated as ‘Benign’, ‘Benign/Likely benign’, ‘Likely benign’ into a single ‘Benign’ category.

Predictions were evaluated using the rank scores downloaded from the database for functional prediction dbNSFP4.2a (Liu et al., 2020) for the following methods: DEOGEN2 (Raimondi et al., 2017), MutationAssessor (Reva et al., 2011), PROVEAN_converted (PROVEAN) (Choi et al., 2012), SIFT_converted (SIFT) (Ng and Henikoff, 2001), SIFT4G_converted (SIFT4G) (Vaser et al., 2016), MutPred (B. Li et al., 2009), MPC (Samocha et al., 2017), LIST-S2 (Malhis et al., 2020), LRT_converted (LRT) (Chun and Fay, 2009), DANN (Quang et al., 2015), MutationTaster_converted (Mutation Taster) (Steinhaus et al., 2021), M-CAP (Jagadeesh et al., 2016), fathmm-MKL_coding (fathmm-MKL) (Shihab et al., 2015), fathmm-XF_coding (fathmm-XF) (Rogers et al., 2018), BayesDel_noAF (BayesDel) (Feng, 2017), REVEL (Ioannidis et al., 2016), CADD_raw (CADD)(Rentzsch et al., 2021), VEST4 (Carter et al., 2013b), MetaLR and MetaSVM (Dong et al., 2015). We also benchmarked five scores that were not in the dbNSFP database, including PrimateAI-3D (Gao et al., 2023), AlphaMissense (Cheng et al., 2023), EVE (Frazer et al., 2021), DeepSequence (Riesselman et al., 2018), and ESM1v (Meier et al., 2021). Due to the unavailability of full mutation effect predictions of the ESM1v model, we reproduced ESM1v scores by applying the ESM1v model from https://github.com/facebookresearch/esm with pretrained-weights to all human protein sequences.

For CDKNA, 37 missense variants were annotated as pathogenic in ClinVar, and 4 missense variants as benign (Table S1). We excluded two pathogenic variants due to insufficient library coverage, and omitted D153Y / c.457G>T based on having a SpliceAI score of 0.76. To expand the set of benign variants, we included variants observed across human populations in gnomAD, TOPMed, and UK Biobank. Since most of these variants were rare (MAF < 0.001), we first compared the performance of our expression and proliferation scores, as well as PrimateAI-3D, across a range of allele frequency thresholds for selecting benign variants (i.e., variants above the threshold), on the task of distinguishing pathogenic variants from benign ones using the area under the receiver operating characteristic curve (AUROC). Since the performance of the four scores was largely consistent across allele frequency thresholds (Figure S2F), and to maximize the set of putatively benign variants, we fixed the threshold at 5e-5, adding 7 variants to the benign set, to comprehensively benchmark a wide range of computational scores (of the 10 variants with allele frequencies > 5e-5, 3 were already in ClinVar and one was excluded due to low coverage in the library). MutationAssessor, LRT, DeepSequence, EVE, and MutPred were excluded from the analysis due to a high number (≥ 27) of missing values for both pathogenic and benign variants.

For TP53, the ClinVar annotation contained 189 pathogenic and 107 benign missense variants (Table S2). One pathogenic variant (G105V) with SpliceAI score > 0.5 was excluded from the analysis. After removing variants that were not scored by every method, the benchmarking set included 182 pathogenic and 66 benign variants. Since no ClinVar variants were annotated in TAD I, we evaluated methods on the task of distinguishing between TAD I variants observed in the population (n = 10) as a proxy for likely benign variants, from those not observed in the population (n = 134) as proxy for likely pathogenic variants. MutPred, DeepSequence, and EVE were excluded from this analysis due to having ≥3 variants with missing scores among the set of 10 likely benign variants.

For SOD1, we evaluated two scores for each model: the raw score and a residualized score obtained by regressing out the proliferation and expression scores from the raw score. Due to the absence of benign variants in ClinVar, we benchmarked scores on the task of distinguishing between 89 ClinVar pathogenic variants and 686 variants that were not classified as pathogenic in other databases or cohorts, as described in the main text. We further excluded variants that were not scored by all methods or that had a SpliceAI score > 0.5, resulting in a common set of 89 pathogenic and 642 benign variants.

To assess statistical differences in AUROCs for each set of benchmarks, we conducted bootstrapping with replacement (sampling 1,000 times) on the shared set of pathogenic and benign/non-pathogenic variants used in each benchmark. Subsequently, Mann-Whitney U tests were used to identify significant differences in AUROCs between pairs of scores.

### TP53 downsampling analysis

To benchmark variant classification performance as a function of total cell numbers and sequencing depth, we randomly sampled the TP53 dataset across a range of mean UMIs per cell (by sampling read counts from a binomial distribution) and cell numbers. We computed an expression score for each downsampling experiment as described in the ‘Expression-derived pathogenicity scores’ section above, with the following exceptions to streamline the analysis: 1) the optimal number of neighbors for UMAP embeddings (10, 15 or 20) and Leiden clustering resolution (ranging from 0.2 to 1 in steps of 0.2) were selected based on the highest concordance, measured by adjusted Rand index, between clusters and TP53 domain annotations; 2) WT samples were not included in training; 3) no sample weighting was applied; and 4) the default set of XGBoost hyperparameters was used for training all downsampled datasets.

For the ClinVar benchmark, we used the common set of variants shared among all downsampling experiments. When calculating Spearman’s correlations between expression and PrimateAI-3D scores, we included all missense variants to account for variants observed in fewer cells, such as variants in TAD I.

### Literature-based annotation of SOD1 enzymatic activity

To test for a correlation between variant effects detected in our assay and effects on SOD1 enzymatic activity, we searched the literature for prior studies that measured the enzymatic activity of individual amino acid variants (Borchelt et al., 1994; Fahmy et al., 2023; Gurney et al., 1994; Hayward et al., 2002; Liochev et al., 1998; Stathopulos et al., 2003). Since different assays were used to measure activity across studies (e.g., pulse radiolysis, gel-based assays, immunoblots), we categorized the enzymatic activity of each variant as normal, reduced, strongly reduced, or ambiguous if there was conflicting evidence (e.g., A5V; Table S4). We found measures of enzymatic activity for 20 variants, two of which had ambiguous effects and were excluded from further analyses.

## Acknowledgements

We thank William Greenleaf, Jay Shendure, J.T. Neal, Lea Starita, Kenneth Matreyek, Douglas Fowler and Fritz Roth for insightful discussions. We additionally thank Eyal Ben-David, Nicole Ersaro, Hong Gao and Jacob Ulirsch for insightful discussions and feedback on the manuscript, and would like to acknowledge the broader Orion Medicines team for their partnership and collaboration.

## Author contributions

Conceptualization: K.K.-H.F., F.A., H.X.

Methodology: F.A., L.C., V.N., K.K.-H.F.

Software: L.C., K.J.-B., F.A.

Validation: L.C., F.A.

Formal Analysis: L.C., K.J.-B., F.A., N.B., J.G., V.N.

Investigation: H.X., M.S., S.W., D.C., X.Z., C.X., T.L., V.Q.

Data Curation: K.J.-B., L.C., F.A.

Writing – Original Draft: F.A.

Writing – Review & Editing: F.A., L.C., K.K.-H.F.

Visualization: F.A., L.C.

Supervision: F.A., H.X., K.K.-H.F., V.N., C.J.Y.

## References

Abel, O., Powell, J.F., Andersen, P.M., Al-Chalabi, A., 2012. ALSoD: A user-friendly online bioinformatics tool for amyotrophic lateral sclerosis genetics. Hum. Mutat. 33, 1345–1351. 10.1002/humu.22157

Adzhubei, I., Jordan, D.M., Sunyaev, S.R., 2013. Predicting functional effect of human missense mutations using PolyPhen-2. Curr. Protoc. Hum. Genet. 76. 10.1002/0471142905.hg0720s76

Attardi, L.D., Reczek, E.E., Cosmas, C., Demicco, E.G., McCurrach, M.E., Lowe, S.W., Jacks, T., 2000. PERP, an apoptosis-associated target of p53, is a novel member of the PMP-22/gas3 family. Genes Dev. 14, 704–718. 10.1101/gad.14.6.704

Bock, C., Datlinger, P., Chardon, F., Coelho, M.A., Dong, M.B., Lawson, K.A., Lu, T., Maroc, L., Norman, T.M., Song, B., Stanley, G., Chen, S., Garnett, M., Li, W., Moffat, J., Qi, L.S., Shapiro, R.S., Shendure, J., Weissman, J.S., Zhuang, X., 2022. High-content CRISPR screening. Nat. Rev. Methods Primers 2. 10.1038/s43586-022-00098-7

Boettcher, S., Miller, P.G., Sharma, R., McConkey, M., Leventhal, M., Krivtsov, A.V., Giacomelli, A.O., Wong, W., Kim, J., Chao, S., Kurppa, K.J., Yang, X., Milenkowic, K., Piccioni, F., Root, D.E., Rücker, F.G., Flamand, Y., Neuberg, D., Lindsley, R.C., Jänne, P.A., Hahn, W.C., Jacks, T., Döhner, H., Armstrong, S.A., Ebert, B.L., 2019. A dominant-negative effect drives selection of *TP53* missense mutations in myeloid malignancies. Science 365, 599–604. 10.1126/science.aax3649

Borchelt, D.R., Lee, M.K., Slunt, H.S., Guarnieri, M., Xu, Z.S., Wong, P.C., Brown, R.H., Jr, Price, D.L., Sisodia, S.S., Cleveland, D.W., 1994. Superoxide dismutase 1 with mutations linked to familial amyotrophic lateral sclerosis possesses significant activity. Proc. Natl. Acad. Sci. U. S. A. 91, 8292–8296. 10.1073/pnas.91.17.8292

Brandes, N., Goldman, G., Wang, C.H., Ye, C.J., Ntranos, V., 2023. Genome-wide prediction of disease variant effects with a deep protein language model. Nat. Genet. 55, 1512–1522. 10.1038/s41588-023-01465-0

Bruijn, L.I., Houseweart, M.K., Kato, S., Anderson, K.L., Anderson, S.D., Ohama, E., Reaume, A.G., Scott, R.W., Cleveland, D.W., 1998. Aggregation and motor neuron toxicity of an ALS-linked SOD1 mutant independent from wild-type SOD1. Science 281, 1851–1854. 10.1126/science.281.5384.1851

Carter, H., Douville, C., Stenson, P.D., Cooper, D.N., Karchin, R., 2013. Identifying Mendelian disease genes with the variant effect scoring tool. BMC Genomics 14 Suppl 3, S3. 10.1186/1471-2164-14-S3-S3

Chen, L.-X., Xu, H.-F., Wang, P.-S., Yang, X.-X., Wu, Z.-Y., Li, H.-F., 2021. SOD1 mutation spectrum and natural history of ALS patients in a 15-year cohort in southeastern China. Front. Genet. 12. 10.3389/fgene.2021.746060

Chène, P., 2003. Inhibiting the p53–MDM2 interaction: an important target for cancer therapy. Nat. Rev. Cancer 3, 102–109. 10.1038/nrc991

Cheng, J., Novati, G., Pan, J., Bycroft, C., Žemgulytė, A., Applebaum, T., Pritzel, A., Wong, L.H., Zielinski, M., Sargeant, T., Schneider, R.G., Senior, A.W., Jumper, J., Hassabis, D., Kohli, P., Avsec, Ž., 2023. Accurate proteome-wide missense variant effect prediction with AlphaMissense. Science 381. 10.1126/science.adg7492

Choi, Y., Sims, G.E., Murphy, S., Miller, J.R., Chan, A.P., 2012. Predicting the functional effect of amino acid substitutions and indels. PLoS One 7, e46688. 10.1371/journal.pone.0046688

Chun, S., Fay, J.C., 2009. Identification of deleterious mutations within three human genomes. Genome Res. 19, 1553–1561. 10.1101/gr.092619.109

Dabrowski, M., Bukowy-Bieryllo, Z., Zietkiewicz, E., 2015. Translational readthrough potential of natural termination codons in eucaryotes--The impact of RNA sequence. RNA Biol. 12, 950–958. 10.1080/15476286.2015.1068497

den Dunnen, J.T., Dalgleish, R., Maglott, D.R., Hart, R.K., Greenblatt, M.S., McGowan-Jordan, J., Roux, A.-F., Smith, T., Antonarakis, S.E., Taschner, P.E.M., on behalf of the Human Genome Variation Society (HGVS), the Human Variome Project (HVP), and the Human Genome Organisation (HUGO), 2016. HGVS recommendations for the description of sequence variants: 2016 update. Hum. Mutat. 37, 564–569. 10.1002/humu.22981

Deng, H.X., Hentati, A., Tainer, J.A., Iqbal, Z., Cayabyab, A., Hung, W.Y., Getzoff, E.D., Hu, P., Herzfeldt, B., Roos, R.P., 1993. Amyotrophic lateral sclerosis and structural defects in Cu,Zn superoxide dismutase. Science. 10.1126/science.8351519

Dong, C., Wei, P., Jian, X., Gibbs, R., Boerwinkle, E., Wang, K., Liu, X., 2015. Comparison and integration of deleteriousness prediction methods for nonsynonymous SNVs in whole exome sequencing studies. Hum. Mol. Genet. 24, 2125–2137. 10.1093/hmg/ddu733

Fahmy, N., Müller, K., Andersen, P.M., Marklund, S.L., Otto, M., Ludolph, A.C., Hamdi, N., 2023. A novel homozygous p.Ser69Pro SOD1 mutation causes severe young-onset ALS with decreased enzyme activity. J. Neurol. 270, 1770–1773. 10.1007/s00415-022-11489-x

Fang, Z., Liu, X., Peltz, G., 2023. GSEApy: a comprehensive package for performing gene set enrichment analysis in Python. Bioinformatics 39. 10.1093/bioinformatics/btac757

Feng, B.-J., 2017. PERCH: A unified framework for disease gene prioritization. Hum. Mutat. 38, 243–251. 10.1002/humu.23158

Findlay, G.M., Daza, R.M., Martin, B., Zhang, M.D., Leith, A.P., Gasperini, M., Janizek, J.D., Huang, X., Starita, L.M., Shendure, J., 2018. Accurate classification of BRCA1 variants with saturation genome editing. Nature 562, 217–222. 10.1038/s41586-018-0461-z

Fischer, N.W., Prodeus, A., Malkin, D., Gariépy, J., 2016. P53 oligomerization status modulates cell fate decisions between growth, arrest and apoptosis. Cell Cycle 15, 3210–3219. 10.1080/15384101.2016.1241917

Fowler, D.M., Fields, S., 2014. Deep mutational scanning: a new style of protein science. Nat. Methods 11, 801–807. 10.1038/nmeth.3027

Frazer, J., Notin, P., Dias, M., Gomez, A., Min, J.K., Brock, K., Gal, Y., Marks, D.S., 2021. Disease variant prediction with deep generative models of evolutionary data. Nature 599, 91–95. 10.1038/s41586-021-04043-8

Gao, H., Hamp, T., Ede, J., Schraiber, J.G., McRae, J., Singer-Berk, M., Yang, Y., Dietrich, A.S.D., Fiziev, P.P., Kuderna, L.F.K., Sundaram, L., Wu, Y., Adhikari, A., Field, Y., Chen, C., Batzoglou, S., Aguet, F., Lemire, G., Reimers, R., Balick, D., Janiak, M.C., Kuhlwilm, M., Orkin, J.D., Manu, S., Valenzuela, A., Bergman, J., Rousselle, M., Silva, F.E., Agueda, L., Blanc, J., Gut, M., de Vries, D., Goodhead, I., Harris, R.A., Raveendran, M., Jensen, A., Chuma, I.S., Horvath, J.E., Hvilsom, C., Juan, D., Frandsen, P., de Melo, F.R., Bertuol, F., Byrne, H., Sampaio, I., Farias, I., do Amaral, J.V., Messias, M., da Silva, M.N.F., Trivedi, M., Rossi, R., Hrbek, T., Andriaholinirina, N., Rabarivola, C.J., Zaramody, A., Jolly, C.J., Phillips-Conroy, J., Wilkerson, G., Abee, C., Simmons, J.H., Fernandez-Duque, E., Kanthaswamy, S., Shiferaw, F., Wu, D., Zhou, L., Shao, Y., Zhang, G., Keyyu, J.D., Knauf, S., Le, M.D., Lizano, E., Merker, S., Navarro, A., Bataillon, T., Nadler, T., Khor, C.C., Lee, J., Tan, P., Lim, W.K., Kitchener, A.C., Zinner, D., Gut, I., Melin, A., Guschanski, K., Schierup, M.H., Beck, R.M.D., Umapathy, G., Roos, C., Boubli, J.P., Lek, M., Sunyaev, S., O’Donnell-Luria, A., Rehm, H.L., Xu, J., Rogers, J., Marques-Bonet, T., Farh, K.K.-H., 2023. The landscape of tolerated genetic variation in humans and primates. Science 380. 10.1126/science.abn8197

Giacomelli, A.O., Yang, X., Lintner, R.E., McFarland, J.M., Duby, M., Kim, J., Howard, T.P., Takeda, D.Y., Ly, S.H., Kim, E., Gannon, H.S., Hurhula, B., Sharpe, T., Goodale, A., Fritchman, B., Steelman, S., Vazquez, F., Tsherniak, A., Aguirre, A.J., Doench, J.G., Piccioni, F., Roberts, C.W.M., Meyerson, M., Getz, G., Johannessen, C.M., Root, D.E., Hahn, W.C., 2018. Mutational processes shape the landscape of TP53 mutations in human cancer. Nat. Genet. 50, 1381–1387. 10.1038/s41588-018-0204-y

Gil, J., Peters, G., 2006. Regulation of the INK4b–ARF–INK4a tumour suppressor locus: all for one or one for all. Nat. Rev. Mol. Cell Biol. 7, 667–677. 10.1038/nrm1987

Goldstein, A.M., Fraser, M.C., Struewing, J.P., Hussussian, C.J., Ranade, K., Zametkin, D.P., Fontaine, L.S., Organic, S.M., Dracopoli, N.C., Clark, W.H., Jr, Tucker, M.A., 1995. Increased risk of pancreatic cancer in melanoma-prone kindreds with*p16*^INK4^mutations. N. Engl. J. Med. 333, 970– 975. 10.1056/nejm199510123331504

Goutman, S.A., Hardiman, O., Al-Chalabi, A., Chió, A., Savelieff, M.G., Kiernan, M.C., Feldman, E.L., 2022. Emerging insights into the complex genetics and pathophysiology of amyotrophic lateral sclerosis. Lancet Neurol. 21, 465–479. 10.1016/S1474-4422(21)00414-2

Gurney, M.E., Pu, H., Chiu, A.Y., Dal Canto, M.C., Polchow, C.Y., Alexander, D.D., Caliendo, J., Hentati, A., Kwon, Y.W., Deng, H.X., 1994. Motor neuron degeneration in mice that express a human Cu,Zn superoxide dismutase mutation. Science 264, 1772–1775. 10.1126/science.8209258

Hafner, A., Bulyk, M.L., Jambhekar, A., Lahav, G., 2019. The multiple mechanisms that regulate p53 activity and cell fate. Nat. Rev. Mol. Cell Biol. 20, 199–210. 10.1038/s41580-019-0110-x

Hayward, L.J., Rodriguez, J.A., Kim, J.W., Tiwari, A., Goto, J.J., Cabelli, D.E., Valentine, J.S., Brown, R.H., Jr, 2002. Decreased metallation and activity in subsets of mutant superoxide dismutases associated with familial amyotrophic lateral sclerosis. J. Biol. Chem. 277, 15923–15931. 10.1074/jbc.M112087200

Hussussian, C.J., Struewing, J.P., Goldstein, A.M., Higgins, P.A., Ally, D.S., Sheahan, M.D., Clark, W.H., Jr, Tucker, M.A., Dracopoli, N.C., 1994. Germline p16 mutations in familial melanoma. Nat. Genet. 8, 15–21. 10.1038/ng0994-15

Ikediobi, O.N., Davies, H., Bignell, G., Edkins, S., Stevens, C., O’Meara, S., Santarius, T., Avis, T., Barthorpe, S., Brackenbury, L., Buck, G., Butler, A., Clements, J., Cole, J., Dicks, E., Forbes, S., Gray, K., Halliday, K., Harrison, R., Hills, K., Hinton, J., Hunter, C., Jenkinson, A., Jones, D., Kosmidou, V., Lugg, R., Menzies, A., Mironenko, T., Parker, A., Perry, J., Raine, K., Richardson, D., Shepherd, R., Small, A., Smith, R., Solomon, H., Stephens, P., Teague, J., Tofts, C., Varian, J., Webb, T., West, S., Widaa, S., Yates, A., Reinhold, W., Weinstein, J.N., Stratton, M.R., Futreal, P.A., Wooster, R., 2006. Mutation analysis of 24 known cancer genes in the NCI-60 cell line set. Mol. Cancer Ther. 5, 2606–2612. 10.1158/1535-7163.mct-06-0433

Ioannidis, N.M., Rothstein, J.H., Pejaver, V., Middha, S., McDonnell, S.K., Baheti, S., Musolf, A., Li, Q., Holzinger, E., Karyadi, D., Cannon-Albright, L.A., Teerlink, C.C., Stanford, J.L., Isaacs, W.B., Xu, J., Cooney, K.A., Lange, E.M., Schleutker, J., Carpten, J.D., Powell, I.J., Cussenot, O., Cancel-Tassin, G., Giles, G.G., MacInnis, R.J., Maier, C., Hsieh, C.-L., Wiklund, F., Catalona, W.J., Foulkes, W.D., Mandal, D., Eeles, R.A., Kote-Jarai, Z., Bustamante, C.D., Schaid, D.J., Hastie, T., Ostrander, E.A., Bailey-Wilson, J.E., Radivojac, P., Thibodeau, S.N., Whittemore, A.S., Sieh, W., 2016. REVEL: An ensemble method for predicting the pathogenicity of rare missense variants. Am. J. Hum. Genet. 99, 877–885. 10.1016/j.ajhg.2016.08.016

Jagadeesh, K.A., Wenger, A.M., Berger, M.J., Guturu, H., Stenson, P.D., Cooper, D.N., Bernstein, J.A., Bejerano, G., 2016. M-CAP eliminates a majority of variants of uncertain significance in clinical exomes at high sensitivity. Nat. Genet. 48, 1581–1586. 10.1038/ng.3703

Jaganathan, K., Kyriazopoulou Panagiotopoulou, S., McRae, J.F., Darbandi, S.F., Knowles, D., Li, Y.I., Kosmicki, J.A., Arbelaez, J., Cui, W., Schwartz, G.B., Chow, E.D., Kanterakis, E., Gao, H., Kia, A., Batzoglou, S., Sanders, S.J., Farh, K.K.-H., 2019. Predicting splicing from primary sequence with deep learning. Cell 176, 535–548.e24. 10.1016/j.cell.2018.12.015

Kakudo, Y., Shibata, H., Otsuka, K., Kato, S., Ishioka, C., 2005. Lack of correlation between p53-dependent transcriptional activity and the ability to induce apoptosis among 179 mutant p53s. Cancer Res. 65, 2108–2114. 10.1158/0008-5472.can-04-2935

Kimura, H., Paranal, R.M., Nanda, N., Wood, L.D., Eshleman, J.R., Hruban, R.H., Goggins, M.G., Klein, A.P., Roberts, N.J., The Familial Pancreatic Cancer Genome Sequencing Project, 2022. Functional CDKN2A assay identifies frequent deleterious alleles misclassified as variants of uncertain significance. Elife 11. 10.7554/elife.71137

Kuleshov, M.V., Jones, M.R., Rouillard, A.D., Fernandez, N.F., Duan, Q., Wang, Z., Koplev, S., Jenkins, S.L., Jagodnik, K.M., Lachmann, A., McDermott, M.G., Monteiro, C.D., Gundersen, G.W., Ma’ayan, A., 2016. Enrichr: a comprehensive gene set enrichment analysis web server 2016 update. Nucleic Acids Res. 44, W90–W97. 10.1093/nar/gkw377

Landrum, M.J., Lee, J.M., Benson, M., Brown, G., Chao, C., Chitipiralla, S., Gu, B., Hart, J., Hoffman, D., Hoover, J., Jang, W., Katz, K., Ovetsky, M., Riley, G., Sethi, A., Tully, R., Villamarin-Salomon, R., Rubinstein, W., Maglott, D.R., 2016. ClinVar: public archive of interpretations of clinically relevant variants. Nucleic Acids Res. 44, D862–D868. 10.1093/nar/gkv1222

Langmead, B., Trapnell, C., Pop, M., Salzberg, S.L., 2009. Ultrafast and memory-efficient alignment of short DNA sequences to the human genome. Genome Biol 10, R25. 10.1186/gb-2009-10-3-r25

Laptenko, O., Tong, D.R., Manfredi, J., Prives, C., 2016. The tail that wags the dog: How the disordered C-terminal domain controls the transcriptional activities of the p53 tumor-suppressor protein. Trends Biochem. Sci. 41, 1022–1034. 10.1016/j.tibs.2016.08.011

Li, B., Krishnan, V.G., Mort, M.E., Xin, F., Kamati, K.K., Cooper, D.N., Mooney, S.D., Radivojac, P., 2009. Automated inference of molecular mechanisms of disease from amino acid substitutions. Bioinformatics 25, 2744–2750. 10.1093/bioinformatics/btp528

Li, H., Handsaker, B., Wysoker, A., Fennell, T., Ruan, J., Homer, N., Marth, G., Abecasis, G., Durbin, R., 1000 Genome Project Data Processing Subgroup, 2009. The Sequence Alignment/Map format and SAMtools. Bioinformatics 25, 2078–2079. 10.1093/bioinformatics/btp352

Lin, J., Chen, J., Elenbaas, B., Levine, A.J., 1994. Several hydrophobic amino acids in the p53 amino-terminal domain are required for transcriptional activation, binding to mdm-2 and the adenovirus 5 E1B 55-kD protein. Genes Dev. 8, 1235–1246. 10.1101/gad.8.10.1235

Liochev, S.I., Chen, L.L., Hallewell, R.A., Fridovich, I., 1998. The familial amyotrophic lateral sclerosis-associated amino acid substitutions E100G, G93A, and G93R do not influence the rate of inactivation of copper- and zinc-containing superoxide dismutase by H2O2. Arch. Biochem. Biophys. 352, 237–239. 10.1006/abbi.1998.0616

Liu, X., Li, C., Mou, C., Dong, Y., Tu, Y., 2020. dbNSFP v4: a comprehensive database of transcript-specific functional predictions and annotations for human nonsynonymous and splice-site SNVs. Genome Med. 12, 103. 10.1186/s13073-020-00803-9

Luecken, M.D., Theis, F.J., 2019. Current best practices in single-cell RNA-seq analysis: a tutorial. Mol. Syst. Biol. 15. 10.15252/msb.20188746

Macosko, E.Z., Basu, A., Satija, R., Nemesh, J., Shekhar, K., Goldman, M., Tirosh, I., Bialas, A.R., Kamitaki, N., Martersteck, E.M., Trombetta, J.J., Weitz, D.A., Sanes, J.R., Shalek, A.K., Regev, A., McCarroll, S.A., 2015. Highly parallel genome-wide expression profiling of individual cells using nanoliter droplets. Cell 161, 1202–1214. 10.1016/j.cell.2015.05.002

Malhis, N., Jacobson, M., Jones, S.J.M., Gsponer, J., 2020. LIST-S2: taxonomy based sorting of deleterious missense mutations across species. Nucleic Acids Res. 48, W154–W161. 10.1093/nar/gkaa288

Matreyek, K.A., Starita, L.M., Stephany, J.J., Martin, B., Chiasson, M.A., Gray, V.E., Kircher, M., Khechaduri, A., Dines, J.N., Hause, R.J., Bhatia, S., Evans, W.E., Relling, M.V., Yang, W., Shendure, J., Fowler, D.M., 2018. Multiplex assessment of protein variant abundance by massively parallel sequencing. Nat. Genet. 50, 874–882. 10.1038/s41588-018-0122-z

Matreyek, K.A., Stephany, J.J., Fowler, D.M., 2017. A platform for functional assessment of large variant libraries in mammalian cells. Nucleic Acids Res. 45, e102. 10.1093/nar/gkx183

McBride, K.A., Ballinger, M.L., Killick, E., Kirk, J., Tattersall, M.H.N., Eeles, R.A., Thomas, D.M., Mitchell, G., 2014. Li-Fraumeni syndrome: cancer risk assessment and clinical management. Nat. Rev. Clin. Oncol. 11, 260–271. 10.1038/nrclinonc.2014.41

McCann, E.P., Henden, L., Fifita, J.A., Zhang, K.Y., Grima, N., Bauer, D.C., Chan Moi Fat, S., Twine, N.A., Pamphlett, R., Kiernan, M.C., Rowe, D.B., Williams, K.L., Blair, I.P., 2021. Evidence for polygenic and oligogenic basis of Australian sporadic amyotrophic lateral sclerosis. J. Med. Genet. 58, 87–95. 10.1136/jmedgenet-2020-106866

McGinnis, C.S., Patterson, D.M., Winkler, J., Conrad, D.N., Hein, M.Y., Srivastava, V., Hu, J.L., Murrow, L.M., Weissman, J.S., Werb, Z., Chow, E.D., Gartner, Z.J., 2019. MULTI-seq: sample multiplexing for single-cell RNA sequencing using lipid-tagged indices. Nat. Methods 16, 619– 626. 10.1038/s41592-019-0433-8

Meier, J., Rao, R., Verkuil, R., Liu, J., Sercu, T., Rives, A., 2021. Language models enable zero-shot prediction of the effects of mutations on protein function. bioRxiv. 10.1101/2021.07.09.450648

Mejzini, R., Flynn, L.L., Pitout, I.L., Fletcher, S., Wilton, S.D., Akkari, P.A., 2019. ALS genetics, mechanisms, and therapeutics: Where are we now? Front. Neurosci. 13, 1310. 10.3389/fnins.2019.01310

Ng, P.C., 2003. SIFT: predicting amino acid changes that affect protein function. Nucleic Acids Res. 31, 3812–3814. 10.1093/nar/gkg509

Ng, P.C., Henikoff, S., 2001. Predicting deleterious amino acid substitutions. Genome Res. 11, 863–874. 10.1101/gr.176601

Nikolova, P.V., Henckel, J., Lane, D.P., Fersht, A.R., 1998. Semirational design of active tumor suppressor p53 DNA binding domain with enhanced stability. Proc. Natl. Acad. Sci. U. S. A. 95, 14675–14680. 10.1073/pnas.95.25.14675

Opie-Martin, S., Iacoangeli, A., Topp, S.D., Abel, O., Mayl, K., Mehta, P.R., Shatunov, A., Fogh, I., Bowles, H., Limbachiya, N., Spargo, T.P., Al-Khleifat, A., Williams, K.L., Jockel-Balsarotti, J., Bali, T., Self, W., Henden, L., Nicholson, G.A., Ticozzi, N., McKenna-Yasek, D., Tang, L., Shaw, P.J., Chio, A., Ludolph, A., Weishaupt, J.H., Landers, J.E., Glass, J.D., Mora, J.S., Robberecht, W., Van Damme, P., McLaughlin, R., Hardiman, O., van den Berg, L., Veldink, J.H., Corcia, P., Stevic, Z., Siddique, N., Silani, V., Blair, I.P., Fan, D.-S., Esselin, F., de la Cruz, E., Camu, W., Basak, N.A., Siddique, T., Miller, T., Brown, R.H., Al-Chalabi, A., Shaw, C.E., 2022. The SOD1-mediated ALS phenotype shows a decoupling between age of symptom onset and disease duration. Nat. Commun. 13, 6901. 10.1038/s41467-022-34620-y

Quang, D., Chen, Y., Xie, X., 2015. DANN: a deep learning approach for annotating the pathogenicity of genetic variants. Bioinformatics 31, 761–763. 10.1093/bioinformatics/btu703

Raimondi, D., Tanyalcin, I., Ferté, J., Gazzo, A., Orlando, G., Lenaerts, T., Rooman, M., Vranken, W., 2017. DEOGEN2: prediction and interactive visualization of single amino acid variant deleteriousness in human proteins. Nucleic Acids Res. 45, W201–W206. 10.1093/nar/gkx390

Raj, N., Attardi, L.D., 2017. The transactivation domains of the p53 protein. Cold Spring Harb. Perspect. Med. 7, a026047. 10.1101/cshperspect.a026047

Rentzsch, P., Schubach, M., Shendure, J., Kircher, M., 2021. CADD-Splice-improving genome-wide variant effect prediction using deep learning-derived splice scores. Genome Med. 13, 31. 10.1186/s13073-021-00835-9

Rentzsch, P., Witten, D., Cooper, G.M., Shendure, J., Kircher, M., 2019. CADD: predicting the deleteriousness of variants throughout the human genome. Nucleic Acids Res. 47, D886–D894. 10.1093/nar/gky1016

Replogle, J.M., Saunders, R.A., Pogson, A.N., Hussmann, J.A., Lenail, A., Guna, A., Mascibroda, L., Wagner, E.J., Adelman, K., Lithwick-Yanai, G., Iremadze, N., Oberstrass, F., Lipson, D., Bonnar, J.L., Jost, M., Norman, T.M., Weissman, J.S., 2022. Mapping information-rich genotype-phenotype landscapes with genome-scale Perturb-seq. Cell 185, 2559–2575.e28. 10.1016/j.cell.2022.05.013

Reva, B., Antipin, Y., Sander, C., 2011. Predicting the functional impact of protein mutations: application to cancer genomics. Nucleic Acids Res. 39, e118. 10.1093/nar/gkr407

Riesselman, A.J., Ingraham, J.B., Marks, D.S., 2018. Deep generative models of genetic variation capture the effects of mutations. Nat. Methods 15, 816–822. 10.1038/s41592-018-0138-4

Rogers, M.F., Shihab, H.A., Mort, M., Cooper, D.N., Gaunt, T.R., Campbell, C., 2018. FATHMM-XF: accurate prediction of pathogenic point mutations via extended features. Bioinformatics 34, 511–513. 10.1093/bioinformatics/btx536

Rosen, D.R., Siddique, T., Patterson, D., Figlewicz, D.A., Sapp, P., Hentati, A., Donaldson, D., Goto, J., O’Regan, J.P., Deng, H.X., 1993. Mutations in Cu/Zn superoxide dismutase gene are associated with familial amyotrophic lateral sclerosis. Nature 362, 59–62. 10.1038/362059a0

Ruffo, P., Perrone, B., Conforti, F.L., 2022. SOD-1 variants in amyotrophic lateral sclerosis: Systematic re-evaluation according to ACMG-AMP guidelines. Genes (Basel) 13, 537. 10.3390/genes13030537

Russo, A.A., Tong, L., Lee, J.O., Jeffrey, P.D., Pavletich, N.P., 1998. Structural basis for inhibition of the cyclin-dependent kinase Cdk6 by the tumour suppressor p16INK4a. Nature 395, 237–243. 10.1038/26155

Rutter, J.L., Goldstein, A.M., Dávila, M.R., Tucker, M.A., Struewing, J.P., 2003. CDKN2A point mutations D153spl(c.457G>T) and IVS2+1G>T result in aberrant splice products affecting both p16INK4a and p14ARF. Oncogene 22, 4444–4448. 10.1038/sj.onc.1206564

Sakaguchi, K., Sakamoto, H., Xie, D., Erickson, J.W., Lewis, M.S., Anderson, C.W., Appella, E., 1997. Effect of phosphorylation on tetramerization of the tumor suppressor protein p53. J. Protein Chem. 16, 553–556. 10.1023/a:1026334116189

Saller, E., 1999. Increased apoptosis induction by 121F mutant p53. EMBO J. 18, 4424–4437. 10.1093/emboj/18.16.4424

Samocha, K.E., Kosmicki, J.A., Karczewski, K.J., O’Donnell-Luria, A.H., Pierce-Hoffman, E., MacArthur, D.G., Neale, B.M., Daly, M.J., 2017. Regional missense constraint improves variant deleteriousness prediction. bioRxiv. 10.1101/148353

Schmidt, R., Steinhart, Z., Layeghi, M., Freimer, J.W., Bueno, R., Nguyen, V.Q., Blaeschke, F., Ye, C.J., Marson, A., 2022. CRISPR activation and interference screens decode stimulation responses in primary human T cells. Science 375. 10.1126/science.abj4008

Shihab, H.A., Rogers, M.F., Gough, J., Mort, M., Cooper, D.N., Day, I.N.M., Gaunt, T.R., Campbell, C., 2015. An integrative approach to predicting the functional effects of non-coding and coding sequence variation. Bioinformatics 31, 1536–1543. 10.1093/bioinformatics/btv009

Stathopulos, P.B., Rumfeldt, J.A.O., Scholz, G.A., Irani, R.A., Frey, H.E., Hallewell, R.A., Lepock, J.R., Meiering, E.M., 2003. Cu/Zn superoxide dismutase mutants associated with amyotrophic lateral sclerosis show enhanced formation of aggregates in vitro. Proc. Natl. Acad. Sci. U. S. A. 100, 7021–7026. 10.1073/pnas.1237797100

Steinhaus, R., Proft, S., Schuelke, M., Cooper, D.N., Schwarz, J.M., Seelow, D., 2021. MutationTaster2021. Nucleic Acids Res. 49, W446–W451. 10.1093/nar/gkab266

Storey, J.D., Tibshirani, R., 2003. Statistical significance for genomewide studies. Proc. Natl. Acad. Sci. U. S. A. 100, 9440–9445. 10.1073/pnas.1530509100

Taylor, J.P., Brown, R.H., Jr, Cleveland, D.W., 2016. Decoding ALS: from genes to mechanism. Nature 539, 197–206. 10.1038/nature20413

Tsherniak, A., Vazquez, F., Montgomery, P.G., Weir, B.A., Kryukov, G., Cowley, G.S., Gill, S., Harrington, W.F., Pantel, S., Krill-Burger, J.M., Meyers, R.M., Ali, L., Goodale, A., Lee, Y., Jiang, G., Hsiao, J., Gerath, W.F.J., Howell, S., Merkel, E., Ghandi, M., Garraway, L.A., Root, D.E., Golub, T.R., Boehm, J.S., Hahn, W.C., 2017. Defining a cancer dependency map. Cell 170, 564–576.e16. 10.1016/j.cell.2017.06.010

Ursu, O., Neal, J.T., Shea, E., Thakore, P.I., Jerby-Arnon, L., Nguyen, L., Dionne, D., Diaz, C., Bauman, J., Mosaad, M.M., Fagre, C., Lo, A., McSharry, M., Giacomelli, A.O., Ly, S.H., Rozenblatt-Rosen, O., Hahn, W.C., Aguirre, A.J., Berger, A.H., Regev, A., Boehm, J.S., 2022. Massively parallel phenotyping of coding variants in cancer with Perturb-seq. Nat. Biotechnol. 40, 896–905. 10.1038/s41587-021-01160-7

Vaser, R., Adusumalli, S., Leng, S.N., Sikic, M., Ng, P.C., 2016. SIFT missense predictions for genomes. Nat. Protoc. 11, 1–9. 10.1038/nprot.2015.123

Wolf, F.A., Angerer, P., Theis, F.J., 2018. SCANPY: large-scale single-cell gene expression data analysis. Genome Biol. 19. 10.1186/s13059-017-1382-0

Yamashita, S., Ando, Y., 2015. Genotype-phenotype relationship in hereditary amyotrophic lateral sclerosis. Transl. Neurodegener. 4. 10.1186/s40035-015-0036-y

